# Treadmilling FtsZ polymers drive the directional movement of sPG-synthesis enzymes *via* a Brownian ratchet mechanism

**DOI:** 10.1101/857813

**Authors:** Joshua W. McCausland, Xinxing Yang, Georgia R. Squyres, Zhixin Lyu, Kevin E. Bruce, Melissa M. Lamanna, Bill Söderström, Ethan C. Garner, Malcolm E. Winkler, Jie Xiao, Jian Liu

**Affiliations:** Department of Biophysics and Biophysical Chemistry, Johns Hopkins School of Medicine, Baltimore, MD, 21205, USA; Department of Molecular and Cellular Biology, Harvard University, Cambridge, MA, 02138, USA; Department of Biology, Indiana University Bloomington, Bloomington, IN, 47405 USA; Structural Cellular Biology Unit, Okinawa Institute of Science and Technology, Japan; and The ithree institute, University of Technology Sydney, Ultimo, NSW, 2007 Australia; Department of Cell Biology, Johns Hopkins School of Medicine, Baltimore, MD, 21205, USA

## Abstract

FtsZ, a highly conserved bacterial tubulin GTPase homolog, is a central component of the cell division machinery in nearly all walled bacteria. FtsZ polymerizes at the future division site and recruits greater than 30 proteins to assemble into a macromolecular complex termed the divisome. Many of these divisome proteins are involved in septal cell wall peptidoglycan (sPG) synthesis. Recent studies found that FtsZ polymers undergo GTP hydrolysis-coupled treadmilling dynamics along the circumference the division site, driving the processive movement of sPG synthesis enzymes. How FtsZ’s treadmilling drives the directional transport of sPG enzymes and what its precise role is in bacterial cell division are unknown. Combining theoretical modeling and experimental testing, we show that FtsZ’s treadmilling drives the directional movement of sPG-synthesis enzymes *via* a Brownian ratchet mechanism, where the shrinking end of FtsZ polymers introduces an asymmetry to rectify diffusions of single sPG enzymes into persistent end-tracking movement. Furthermore, we show that the processivity of this directional movement is dependent on the binding potential between FtsZ and the enzyme, and hinges on the balance between the enzyme’s diffusion and FtsZ’s treadmilling speed. This interplay could provide a mechanism to control the level of available enzymes for active sPG synthesis both in time and space, explaining the distinct roles of FtsZ treadmilling in modulating cell wall constriction rate observed in different bacterial species.

---

During cell wall constriction in most Gram-negative bacteria, new septal peptidoglycan (sPG) synthesis and old cell wall degradation occur simultaneously^1^. A large number of the cell wall enzymes involved in this process and their regulators have been identified. However, it remains unclear how these proteins are orchestrated in time and space to achieve successful cytokinesis and at the same time maintain the structural integrity of the septal cell wall^2,3^. Perturbations of PG remodeling at septum compromise cell division and often lead to cell lysis^4^.

Recent studies have indicated that FtsZ, an essential component of the bacterial cell division machinery, may play a central part in regulating the spatiotemporal coordination of sPG synthesis enzymes. FtsZ is a highly conserved bacterial tubulin homologue and GTPase^5-7^. During cell division, FtsZ polymerizes at the cytoplasmic face of the inner membrane to form a ring-like structure (Z-ring) at mid-cell^8-10^. The Z-ring then locally recruits an ensemble of more than 30 proteins, many of which are sPG-remodeling enzymes^1,11^, to initiate septal cell wall constriction. New studies employing super-resolution and single-molecule imaging *in vitro* and *in vivo* have demonstrated that the FtsZ polymers exhibit GTP hydrolysis-driven treadmilling dynamics, which are the continuous polymerization at one end and depolymerization at the other end, with individual FtsZ monomers remaining stationary in the middle^12-15^. Most interestingly, it was found that FtsZ’s treadmilling dynamics drive processive movements of the essential sPG transpeptidase (TPase, FtsI in *E*. *coli* and PBP2B in *B*. *subtilis*)^12,13^ and glycosyltransferase FtsW^16^. Consequently, it was proposed that FtsZ’s treadmilling dynamics spatially and temporally distribute sPG synthesis enzymes along the septum plane to ensure smooth septum morphogenesis^13^. However, it is unknown how FtsZ’s treadmilling dynamics with stationary monomers in the cytoplasm are transduced into the periplasm to drive the persistent and directional movement of cell wall synthesis enzymes. The role of FtsZ’s treadmilling dynamics in modulating sPG synthesis activity also remains elusive, as it was shown that the cell wall constriction rate is dependent on FtsZ’s treadmilling speed in *B*. *subtilis*^12^ but not in *E*. *coli* ^13^, or *S*. *aureus*^*17*^.

In this work, we combined agent-based theoretical modeling with single-molecule imaging-based experimental testing to address the mechanism of the FtsZ treadmilling-dependent processive movement of sPG enzymes and its associated role in bacterial cell division. We found that a Brownian ratchet mechanism underlies the persistent and directional movement of single sPG synthesis enzyme molecules driven by FtsZ’s treadmilling dynamics. Using FtsI as a model sPG enzyme, we found that the processivity of the Brownian ratchet is dependent on the (indirect) binding potential between FtsI and FtsZ and modulated by the balance between FtsI’s random diffusion and FtsZ’s treadmilling speed. This finding offers predictions about how different bacterial species could harness the same FtsZ treadmilling machinery to achieve distinct processivities of sPG enzymes so that the available level of sPG synthases for cell wall constriction can be controlled differentially. Given the lack of linear stepping motors in the prokaryotic world, our work suggests a general framework for how polymer dynamics coupled with Brownian ratcheting could underlie directional transport of cargos and be shaped by evolution to meet the needs of different cellular milieus.

## Results

### Model description

Our model is based on the concept of a Brownian ratchet, where FtsZ’s treadmilling introduces an asymmetry to bias the random diffusion of FtsI molecules in the periplasm, upon which FtsI persistently follows the shrinking end of a treadmilling FtsZ filament (Fig. 1). The quantitative details of the model are rooted in the physical and chemical properties of key components of the system, which can be characterized by experiments.

**Figure 1.**
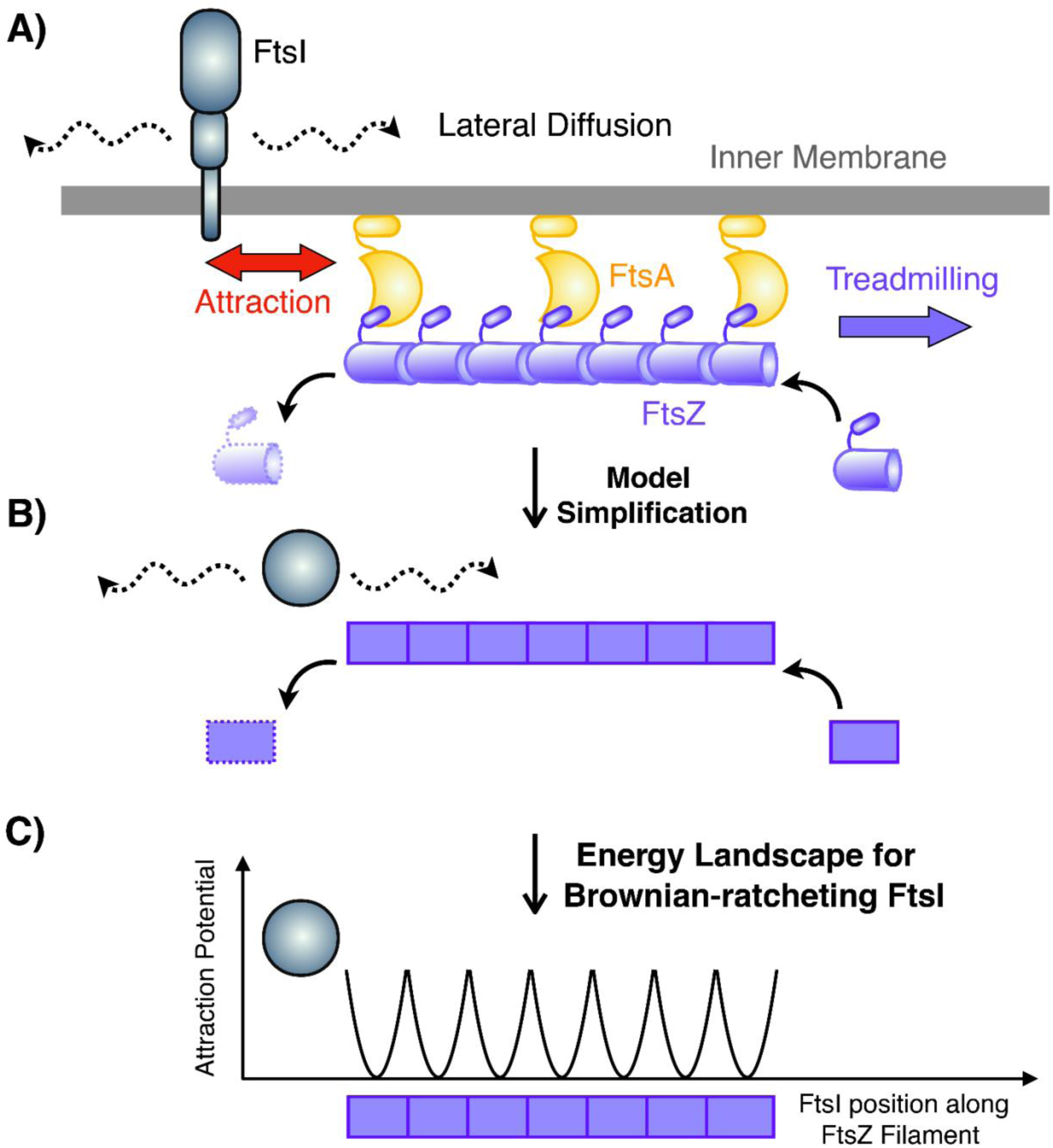
Model description. A) Schematic representation of sPG synthase complex’s interaction with FtsZ treadmilling. FtsZ resides in the cytoplasm. FtsI is a transmembrane protein that does not dissociate from the membrane even when it dissociates from FstZ. B) Model simplification of FtsZ – FtsI interaction at the septum. The FtsZ filament (purple) undergoes treadmilling by dissociating FtsZ subunit from the left end and associating new ones from the right end. While the FtsI complex (grey) intrinsically diffuses around, it has a binding affinity to FtsZ subunits. C) Schematics of FtsZ – FtsI binding potentials. Here, the binding potential is assumed to be harmonic and short-ranged (∼ 5 nm), which is about the size of an individual FtsZ subunit. Note that there is no energy barrier for FtsI to bind to FtsZ. Once the FtsI binds to a FtsZ subunit, however, the binding potential presents an energetic barrier preventing the FtsI from diffusing away.

As shown in Fig. 1, the model describes the movement of a free FtsI molecule at the septum as quasi-1D. The model assumes that FtsI, an essential TPase with a single transmembrane domain and a cytoplasmic tail, can freely diffuse along the inner membrane at the septum or interact indirectly with a treadmilling FtsZ filament underneath the inner membrane (Fig. 1A). The dynamics of a single FtsI molecule at the septum are therefore determined by three parameters: the constant of FtsI’s free diffusion (*D*), the treadmilling speed (*V*_Z_) of FtsZ filaments, and the attraction force determined by the binding potential (*U*) between FtsI and FtsZ (Fig. 1B and C).

To set the ranges of the three parameters, we consider the following. First, we set the diffusion constant to range from 10^−3^ to 10^−1^ μm^2^/s, which is of a typical inner membrane protein in bacterial cells^18,19^. For example, PBP2, the counterpart of FtsI in cell wall elongation, was measured at ∼ 0.06 μm^2^/s^18,19^. Second, the average treadmilling speed of FtsZ was at ∼ 20 – 40 nm/s *in vivo* but can be a few-fold faster *in vitro*, therefore we set a large range of 10 – 100 nm/s^14,15^. Third, FtsI interacts with FtsZ at the septum indirectly through a relay of protein-protein interactions that include FtsN, FtsA, and/or FtsEX^1,20^. For simplicity we omit the details of the protein-protein interaction relay and refer to it as the interaction between FtsI and FtsZ. The indirect interaction between an FtsZ monomer and a nearby FtsI molecule constitutes an attractive force for each other and can be described as a short-ranged harmonic binding potential (Fig. 1C). We assumed the potential range was ± 2.5 nm, commensurate with the size of an FtsZ monomer (∼ 5 nm)^21-24^. The potential’s magnitude was ∼10 *k*_B_T, corresponding to a *K*_d_ in the μM range, which is typical for protein-protein interactions in the bacterial divisome system^25-31^. Note that here we use FtsI in *E*. *coli* as the model sPG enzyme, but the same analysis can be applied to any other sPG enzyme or divisome proteins as well.

To numerically compute the model, we describe the dynamics of FtsI by a Langevin-type equation (Equation (1)) where the viscous drag force on the molecule is in balance with a driving force *f* and a force from thermal noise *ξ*:

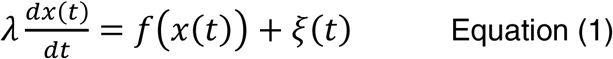

Here, *x*(*t*) represents the location of a FtsI molecule at time *t* along the 1D septum. *λ* is the effective viscous drag coefficient for FtsI’s movement with *λ* = *k*_B_T/*D*, where *D* is the diffusion constant of free FtsI molecules on the inner membrane when it is not interacting with FtsZ. *f*(*x*(*t*)) is the attractive force exerted upon the FtsI molecule by the FtsZ’s binding potential, *U*(*x, t*) (Fig. 1C). Specifically, *f*(*x*(*t*)) = – ∂*U*(*x, t*)/∂*x* at time *t*. The last term *ξ*(*t*) reflects the random diffusive motion of the FtsI on the inner membrane with ⟨*ξ*(*t*)·*ξ*(*t′*)⟩ = 2*D*·Δ*t*·δ(*t*–*t′*), where Δ*t* is the unitary time step in simulation.

Next, we simulate the treadmilling of an FtsZ filament. The model depicts the filament shrinking and growing according to Equations (2) and (3), which respectively describe how the positions of the shrinking and growing ends of a treadmilling FtsZ filament at time t, x_*S*_(*t*) and x_*G*_(*t*) are related to the treadmilling rate, *V*_*Z*_:

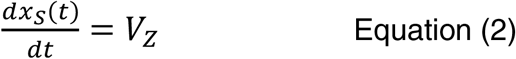

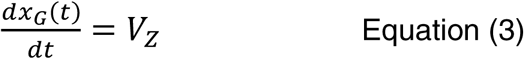

The model simulates the treadmilling events in a discretized sense, which occur every 5nm/(*Vz*·Δ*t*) simulation time step. Each time an FtsZ subunit falls off the shrinking end of the filament, the associated binding potential vanishes with it; likewise, when an FtsZ subunit adds onto the growing end, the associated binding potential appears with it. The treadmilling speed *V*z of each FtsZ filament is drawn stochastically from an experimentally measured distribution^13^. The FtsZ filament length is set at 50 monomers (250 nm) and the treadmilling speed is independent of the filament length, in accordance to previous biochemical studies and a recent *in vivo* study^20,32-35^. To discern principal interactions, the model considers one FtsI molecule and one FtsZ filament in a self-contained septal section. It can be easily expanded to include multiple FtsI molecules per FtsZ filament (Fig. S1).

Assuming that the FtsZ filament treadmills from left to right with a steady-state length of ∼ 250 nm, the model implements open boundary conditions on the FtsI molecule at both the left and right edges of the system and the right-ward FtsZ treadmilling is not limited. The model results presented below reflect the nominal case, whose essence remains robust against variations of model parameters within the physical range constrained by existing experimental data.

### A Brownian ratchet mechanism couples FtsI’s directional movement to FtsZ’s treadmilling dynamics at the shrinking end

As we described above, Brownian ratcheting hinges on the diffusion of FtsI, its interaction with FtsZ, and FtsZ’s treadmilling speed. To examine how the movement of FtsI depends on FtsI’s diffusion and the binding potential between FtsI and FtsZ, we kept FtsZ’s treadmilling speed constant at an experimentally measured speed of 25 nm/s and carried out a phase diagram study using stochastic simulations. As described above, we chose a parameter range of 0.0001 to 0.1 μm^2^/s for FtsI’s diffusion^18,19^. The upper limit of the binding potential was set ∼ 20 *k*_B_T, which corresponded to a dissociation constant *K*_d_ in the nM-range.

We considered an initial condition in which both the shrinking end of an FtsZ filament and an FtsI molecule were at the left boundary of the septal section. To be commensurate with our experimental analysis, we counted an FtsI trajectory as moving directionally if it tracked the shrinking end of a treadmilling FtsZ filament persistently and unidirectionally for at least 4 seconds. Because of the stochastic nature of Brownian ratcheting, we characterized the state of FtsI under this parameter set condition as persistent end-tracking in the phase diagram if 50% or more of simulated FtsI trajectories displayed such a persistent directional movement.

As shown in the phase diagram in Fig. 2A, the model showed that when the binding potential between FtsZ and FtsI was weak (< 5 *k*_B_T, ∼ mM Kd), FtsI largely displayed random diffusion without directional movements along the septum. When the attraction potential was sufficiently strong (> 5 *k*_B_T), strong binding quenched free diffusion and confined FtsI to the end of an FtsZ filament. As the FtsZ subunit at the shrinking end of the filament fell off, the next one in the row attracted and coupled to FtsI, which pulled FtsI to the right by ∼ 5 nm. With the subsequent FtsZ subunits falling off one after the other from the shrinking end, the FtsI molecule ratcheted forward and persistently tracked the end of the treadmilling FtsZ filament. These consecutive movements resulted in a persistent and directional trajectory of FtsI (Fig. 2B). Consequently, the speed of FtsI directional movement was tightly coupled to FtsZ’s treadmilling speed (Fig. 2C), recapitulating the experimentally measured near-linear correlation between FtsI’s directional motion with FtsZ’s treadmilling speeds in both wildtype and FtsZ GTPase mutants^13^.

**Figure 2.**
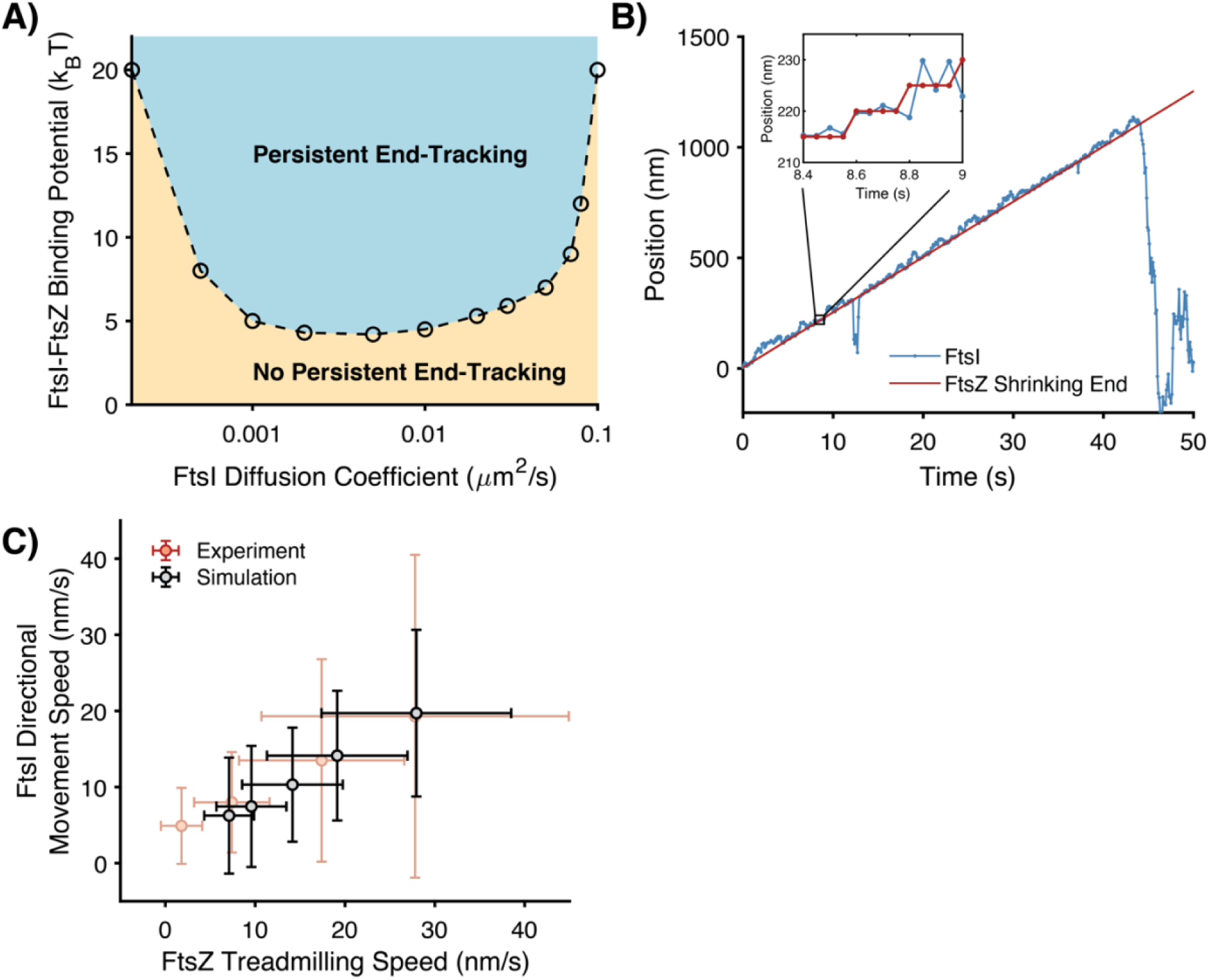
A FtsZ treadmilling-mediated Brownian ratchet mechanism drives FtsI’s directional movement. A) Phase diagram depicting the dependence of FtsI’s persistent end-tracking (blue-shaded region) on FtsI’s diffusion constant and FtsI-FtsZ binding potential. B) A representative simulated trajectory of a FtsI molecule persistently end-tracking with a treadmilling FtsZ filament. Inset: zoom-in view of a boxed region of the trajectory. The model parameters for this simulation are: FtsZ’s treadmilling speed *V*_Z_ = 25 nm/s, FtsI diffusion constant *D* = 0.04 μm^2^/s, FtsZ–FtsI binding potential *U* = 10 *k*_B_T, and the simulation time step = 5×10^−6^ s. The full trajectory is plotted every 10^−1^ s, and the zoom-in inset is plotted every 5×10^−2^ s. C) FtsI’s directional speed tightly couples with FtsZ’s treadmilling speed. Each of the data points was the average of >80 independent stochastic simulation trajectories using the segments that undergo directional movement. FtsZ’s treadmilling speed was fixed within each trajectory but varied across the ensemble following a Gaussian distribution with a standard deviation (SD) of 30%, in line with the experimental measurements in FtsZ WT and GTPase mutants^13,16^.

The phase diagram (Fig. 2A) also showed that at a constant binding potential between FtsZ and FtsI, persistent end-tracking of FtsI required an appropriate range of diffusion constants. If FtsI diffused too rapidly, it could not be confined by the binding potential of the shrinking end of the FtsZ filament. Conversely, when FtsI diffused too slowly, it was not able to keep up with the speed of departing FtsZ subunits at the shrinking end. Once it fell behind, the FtsI molecule lost contact with the left most FtsZ subunits permanently.

We note that an alternative initial condition, in which an FtsI molecule binds in the middle of a FtsZ filament, will result in the same end-tracking behavior. The bound FtsI molecule, if has not dissociated, will start end-tracking when the shrinking end of the FtsZ polymer approaches and mobilizes it (Fig. S1).

Our modeling also found that coupling to the growing end of an FtsZ filament is unable to produce the directional movement of FtsI molecules (Fig. S2). As there is no biochemical evidence showing that the FtsI-binding potential of a newly added FtsZ subunit at the growing tip will be higher than the ones in the middle of the filament, the addition of a new FtsZ subunit at the growing end does not bias the diffusion of the FtsI molecule bound at the original tip to dissociate and re-associate with the new FtsZ subunit. Consequently, slow-diffusing FtsI molecules will be stuck in the local binding potential, unable to catch up with the addition of new FtsZ subunits (Fig. S2A), whereas fast-diffusing FtsI molecules have a high probability of escaping from the tip, because there are no FtsZ subunits beyond the growing tip to keep it within the vicinity as that in the shrinking tip-tracking scenario (Fig. S2B). Therefore, FtsI cannot persistently track the growing end of a FtsZ filament.

Taken together, our analysis showed that the end-tracking Brownian ratchet mechanism was able to couple FtsI’s directional movement to FtsZ’s shrinking end within the parameter range that is well consistent with experimentally measured data. Furthermore, the same model could explain the nondirectional movement of the cytoplasmic tail of FtsN, another divisome protein, in a recent *in vitro* study^25^. In this study, the cytoplasmic tail of FtsN was reported to follow the tracks of treadmilling FtsZ filaments on a supported lipid bilayer at the ensemble level. At the single molecule level, however, the FtsN tail only binds and unbinds FtsZ filaments transiently but does not exhibit directional movement^25^. Such a scenario could be explained by to our Brownian ratchet model in that the diffusion of free FtsN cytoplasmic tail anchored on the membrane was too large (0.3 – 0.6 µm^2^/s)^25^.

### FtsZ’s treadmilling speed modulates processivity of FtsI’s end-tracking

Next, we investigated how FtsZ’s treadmilling speed impacts the processivity of FtsI’s directional movement at the shrinking end. Addressing this question will help us understand the role of FtsZ’s treadmilling dynamics in the spatial organization and/or regulation of sPG synthesis activity. We focused on three features that collectively define the processivity of FtsI’s end-tracking: (1) the propensity, (2) the run distance, and (3) the duration time of persistent end-tracking trajectories.

We first examined how the relative propensity of FtsI’s persistent end-tracking was modulated by FtsZ treadmilling speed. The relative propensity is defined as the percentage of the number of FtsI persistent end-tracking trajectories at each FtsZ treadmilling speed, normalized by the total number of FtsI persistent end-tracking trajectories of all the simulated FtsZ treadmilling speeds. Keeping the diffusion constant of FtsI at 0.04 μm^2^/s and the binding potential at 10 *k*_B_T, stochastic simulations of the Brownian ratchet model predicted that the relative propensity of persistent end-tracking trajectories of FtsI dropped off with increasing FtsZ’s treadmilling speed (Fig. 3A). That is, when FtsZ treadmills too fast, FtsI could not persistently track the FtsZ shrinking end in most cases and became largely diffusive.

**Figure 3.**
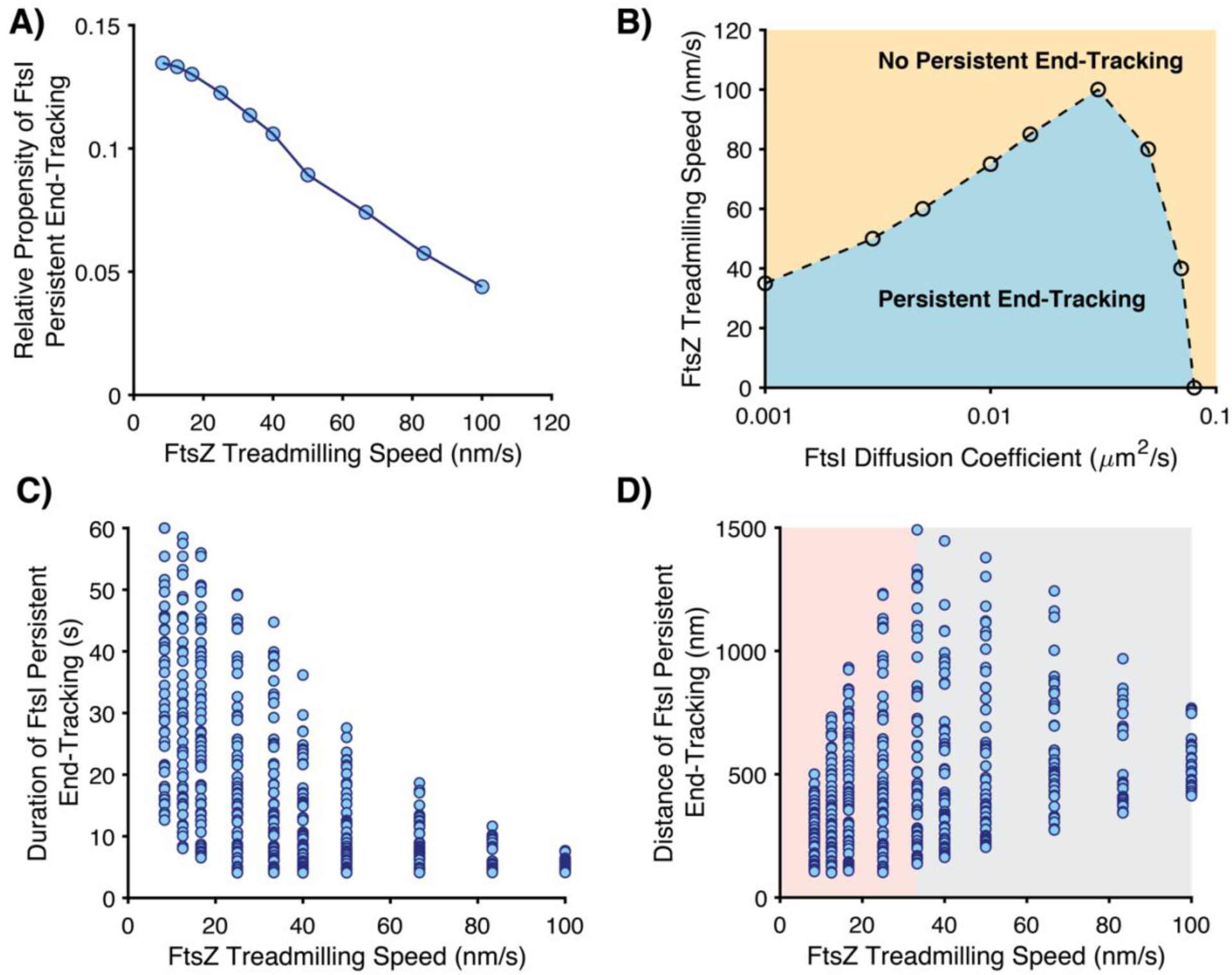
Model predictions for the processivity of FtsI directional movement modulated by FtsZ treadmilling speed. A) Predicted relative propensity of FtsI persistent end-tracking as a function of FtsZ’s treadmilling speed. For each FtsZ’s treadmilling speed, 100 independent stochastic FtsI trajectories were simulated, from which the number of FtsI persistent end-tracking trajectories were counted. In line with our experimental measurement, the criteria for persistent end-tracking were as follows: (1) the distance between the FtsZ shrinking end and the FtsI is less than 100 nm and (2) FtsI persistently follows the FtsZ shrinking end for greater than 4 seconds. A total of 664 out of 1000 simulated FtsI trajectories were scored as persistent end-tracking, and the number of the trajectories at each FtsZ’s treadmilling speed was normalized as a relative probability of end-tracking. B) Calculated phase diagram of FtsI persistent end-tracking characterized by FtsI diffusion constant and FtsZ treadmilling speed. C) Predicted FtsZ treadmilling speed-dependence of the duration of FtsI persistent end-tracking. D) Predicted FtsZ treadmilling speed-dependence of run distance of FtsI persistent end-tracking. For the model calculations in (A-D), the FtsZ–FtsI binding potential is set to be 10 *k*_B_T.

To further this point, we calculated the phase diagram of FtsI’s persistent end-tracking propensity as a function of both FtsI’s diffusion constant and FtsZ’s treadmilling speed (Fig. 3B), while keeping the binding potential fixed at 10 *k*_B_T. Again, we used a threshold of 50% FtsI persistent end-tracking trajectories as the criterion for the phase boundary. As shown in Fig. 3B, for a fixed diffusion constant of FtsI, there was an upper limit of FtsZ’s treadmilling speed that FtsI could persistently track. Conversely, for a fixed FtsZ treadmilling speed, persistent end-tracking of FtsI required an appropriate range of diffusion constants. Importantly, very large diffusion constants of FtsI (> 0.1 μm^2^/s) did not support persistent end-tracking irrespective of FtsZ’s treadmilling speed. These results were consistent with the phase diagram in Fig. 2A and again the recent *in vitro* study of FtsN’s cytoplasmic tail^25^.

Next, we investigated how FtsZ’s treadmilling speed modulates the run distance and duration time of FtsI’s persistent end-tracking. The Brownian ratchet model predicted that both the run length and duration time of FtsI’s persistent end-tracking should display broad distributions due to the stochastic nature of FtsI’s diffusion and the interaction between FtsI and FtsZ. Moreover, the model predicts that when FtsZ’s treadmilling speed increases, the duration time of FtsI’s persistent end-tracking will decrease (Fig. 3C), whereas the run distance will display a biphasic dependence – it increases to peak around an intermediate FtsZ’s treadmilling speed (∼ 30 nm/s at the current parameter setting), and then decreases when FtsZ’s treadmilling speed increases further (Fig. 3D). Importantly, such distinctive dependences of duration time and run distance on FtsZ’s treadmilling speed is a natural consequence of the Brownian ratchet mechanism (SI).

Qualitatively speaking, when an FtsZ subunit falls off to the cytoplasm from the shrinking end of the FtsZ filament, the associated FtsI molecule will dissociate from the FtsZ subunit, either diffuse away on the membrane, or catch up with the next FtsZ subunit in the row to continue end-tracking, the latter depending on how fast FtsZ treadmills. When FtsZ treadmills too fast (for example > 30 nm/s), it will be difficult for FtsI to catch up (Fig. 3A), resulting in early termination of end-tracking, and hence both the persistence run distance and duration time will be short (right sides of Fig. 3C and D). When FtsZ treadmills relatively slowly (< 30 nm/s), the probability of FtsI catching up with the shrinking end of the FtsZ filament is high (Fig. 3A). Therefore, the slower FtsZ treadmills, the fewer number of dissociation events of an end-tracking FtsI molecule would face, and hence the lower the chance for FtsI to diffuse away, leading to a longer time duration of persistence run (Fig. 3C). Within the same time window, however, the persistence run distance will be proportional to the FtsZ’s treadmilling speed as predicted in Fig. 3D, that is, the slower FtsZ treadmills, the shorter FtsI’s persistence run distance is. One can imagine in the extreme case where FtsZ does not treadmill at all (*V*_Z_ = 0), the duration time of persistence runs would then be the longest and mainly dictated by the intrinsic dissociation rate of FtsI from FtsZ, and the persistence run distance would be the shortest (i.e., the size of a single FtsZ subunit). An analytical proof of these relationships is provided in the SI and Fig. S3.

### Single-molecule tracking of FtsI confirms model predictions

To experimentally examine the model’s predictions on the modulation of the processivity of FtsI’s directional movement by FtsZ’s treadmilling speed, we performed single-molecule tracking (SMT) of a functional sandwich fusion protein Halo-FtsI_SW_ labeled with JF646 in live *E*. *coli* cells^36,37^. To avoid disrupting the cytoplasmic interactions of FtsI’s N-terminal tail with other divisome proteins, we inserted the Halo tag between the last residue (18) of the N-terminal cytoplasmic tail and the first residue of the inner membrane helix (19) of FtsI (Fig. 4A). We integrated the *halo-ftsI*_*SW*_ fusion gene into the chromosome replacing the endogenous *ftsI* gene and showed that it was expressed as a full-length fusion protein and supported normal cell division as a sole cellular source of FtsI similar to wild-type (WT) cells (Fig. S4).

**Figure 4.**
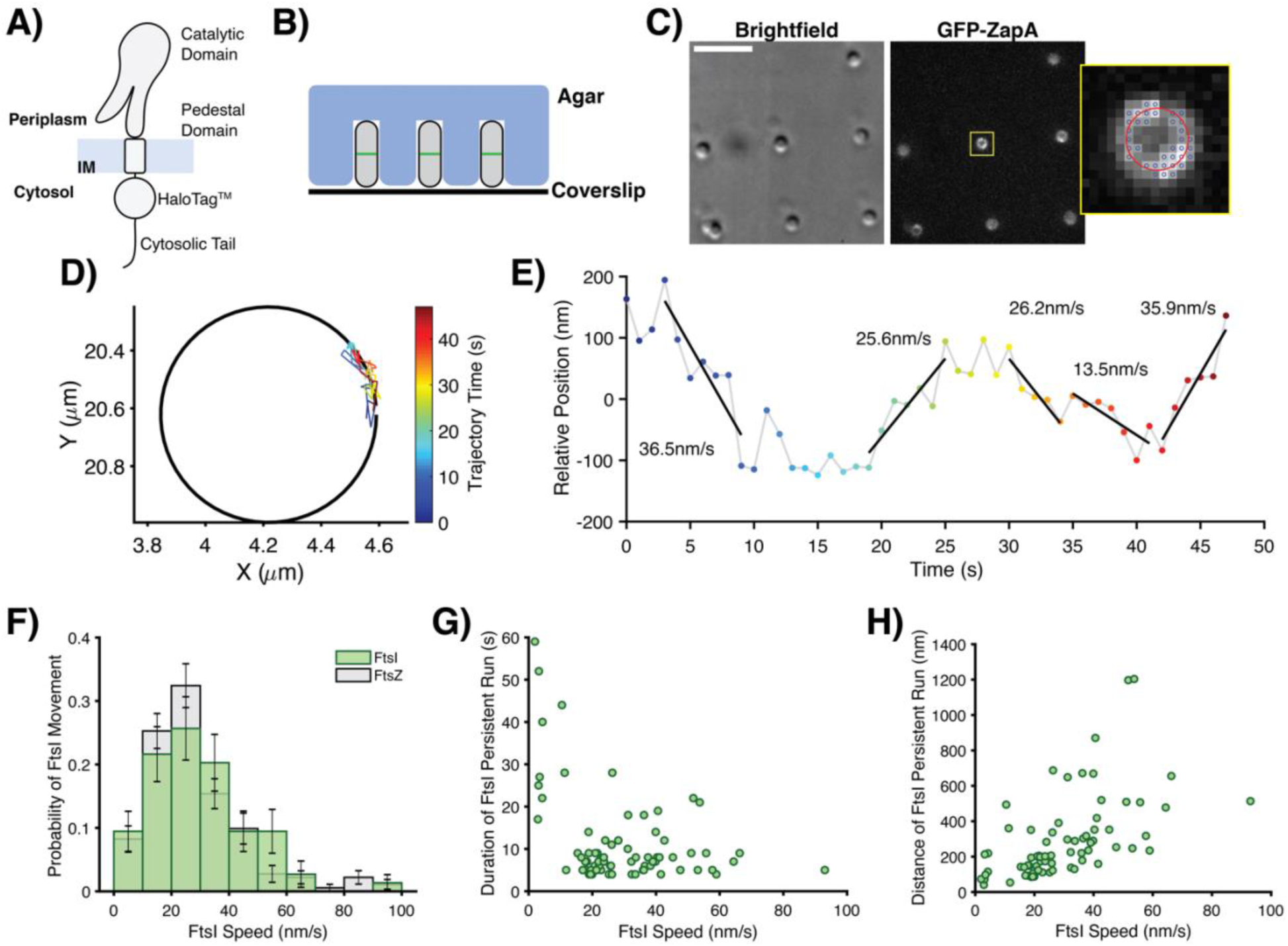
Experimental characterization of FtsI directional motion. A) Schematic of the functional sandwich fusion of FtsI. The Halo tag is inserted between residue 18 and 19 of FtsI, immediately before the first residue of the TM domain. B) Diagram of individual *E*. *coli* cells loaded in microholes made by nanopillars. C) Brightfield and fluorescence images of microholes loaded with *E*. *coli* cells labeled with GFP-ZapA and Halo-FtsI_SW_ fusion proteins. Inset shows the zoomed image of one cell in the yellow box. Blue circles indicated the pixels used for the fit of ZapA-GFP circle. D) Circle-fitting of GFP-ZapA image super-imposed with the trajectory of a single FtsI molecule, colored in time. E) the unwrapped trajectory from D with fitted lines at each segment to extract directional speeds, persistent run duration and distance. F) Histogram of FtsI’s directional movement speeds (N = 77 trajectories, 74 directional events), with the distribution of FtsZ’s treadmilling speed presented in grey for reference (from Yang et al.^13^). For the purpose of this study, we only plotted the fast-moving population of FtsI that follows FtsZ’s treadmilling and leave out the slow-moving population of FtsI that we show to be independent of FtsZ’s treadmilling^13^. Both FtsI’s and FtsZ’s histograms were bootstrapped 100 times for error bars (showing standard error) and a KS test showed no significant difference between the two (p = 0.1852). G) Dependence of the duration of FtsI’s persistent run on its speed. H) Dependence of the distance of FtsI’s persistent run on its speed.

To obtain precise measurements of the persistent run distance and duration time of single Halo-FtsI_SW_ molecules, we trapped individual *E*. *coli* cells vertically in agarose microholes made using cell-shaped nanopillar molds as previously described^38,39^ so that the entire circumference of the septum could be visualized at the same focal plane (Fig. 4B). To determine whether a Halo-FtsI_SW_ molecule was at a septum, we labeled the FtsZ-ring using an ectopically expressed GFP-ZapA fusion protein, which we and others have previously shown as a faithful marker of the Z-ring localization and dynamics^40^. The GFP-ZapA image also allowed us to unwrap the circular trajectories of FtsI-Halo molecules along the septum (Fig. 4D) to linear displacements along the circumference of the septum (Fig. 4E), from which we could measure the persistent run speed, distance, and duration time (Fig. 4F, G and H).

As shown in Fig. 4F, the directional motion speed of Halo-FtsI exhibited a wide distribution, similar to what we previously observed for FtsZ’s treadmilling. The similarity between FtsI’s directional motion speed distribution and FtsZ’s treadmilling speed distribution suggest that at these speed ranges, *E*. *coli* FtsI can faithfully end-track treadmilling FtsZ filaments as the model predicted, a point that will become important in the section below. Most importantly, the persistence run distance and duration time exhibited largely the same trends as what were predicted by the model: while the run duration time decreased monotonically (Fig. 4G), the persistence run distance increased and then decreased when FtsI’s speed increased (Fig. 4H). Note here that we inferred FtsZ’s treadmilling speed from FtsI’s directional motion speed due to the difficulty of a two-color co-tracking experiment in the same cells and because we have demonstrated previously that these two were linearly coupled^13^. Another potential caveat in these experiments was that a very fast FtsZ’s treadmilling speed (*i*.*e*., > 80 nm/s) is rare in wildtype *E*. *coli* cells as we showed previously. Therefore, given the relatively small dataset for high speed FtsZ treadmilling, our data cannot definitively determine whether FtsI could effectively end-track FtsZ filaments of very fast treadmilling speeds. Nevertheless, the agreement of our experimental measurements with theoretical predictions supported the validity of the Brownian ratchet model.

### sPG synthase’s diffusion and FtsZ-binding potential underlie the dependence of its enzymatic activity on FtsZ treadmilling

In *E*. *coli*, the total amount of septal PG synthesis and the septum constriction rate are insensitive to perturbations in FtsZ’s treadmilling speed from ∼ 8 nm/s to ∼ 30 nm/s in a series of FtsZ GTPase mutants^13^. This insensitivity suggests that end-tracking, directionally-moving FtsI molecules were inactive in sPG synthesis. Indeed, a second, slow-moving population of FtsW and FtsI (∼ 8 nm/s) is found to move independently of FtsZ’s treadmilling, and likely corresponds to the active population of SPG synthesis in *E*. *coli*^16^. Similarly, in. *S*. *pneumoniae*, FtsW and its cognate TPase PBP2x were found to move completely independently of FtsZ’s treadmilling^41^, likely representing the active population of sPG synthase as that in *E*. *coli*. In *B*. *subtilis*, however, it was shown that the Z-ring constriction speed is positively correlated with FtsZ’s treadmilling speed, suggesting that the faster FtsZ treadmills, the higher the sPG synthesis activity^12^. How could the same FtsZ treadmilling dynamics result in different sPG synthesis activity in different species?

We propose that since FtsI molecules tracking FtsZ filaments are most likely inactive^16^, the population of FtsI molecules not tracking with treadmilling FtsZ polymers is then available for sPG synthesis. Therefore, FtsI’s off-rate, or the reciprocal of the time a FtsI molecule spends bound in the middle of FtsZ polymers and/or persistently end-tracking, represents the rate at which an FtsI molecule becomes available for sPG synthesis, and therefore is proportional to the sPG synthesis rate. As such, FtsZ’s treadmilling speed could modulate the rate of sPG synthesis in different bacterial species depending on the unique combination of the enzyme’s diffusion coefficient and binding potentials. This modulation could explain the difference observed between *E*. *coli* and *B*. *subtilis*.

As shown in Fig. 5A, the model suggests that when FtsI diffuses relatively fast (∼ 0.05 μm^2^/s, blue line), the lifetime of FtsZ-bound FtsI is largely insensitive to FtsZ’s treadmilling speed in a 3-fold range from ∼ 8 nm/s to 25 nm/s. In contrast, when FtsI diffuses relatively slowly, the lifetime of FtsZ-bound FtsI is critically dependent on FtsZ treadmilling speed. For example, at a diffusion constant of 0.005 μm^2^/s, the relative lifetime of FtsI molecules decreased by ∼ 70% when FtsZ treadmilling speed increased from ∼ 8 nm/s to 25 nm/s (Fig. 5A, green line).

**Figure 5.**
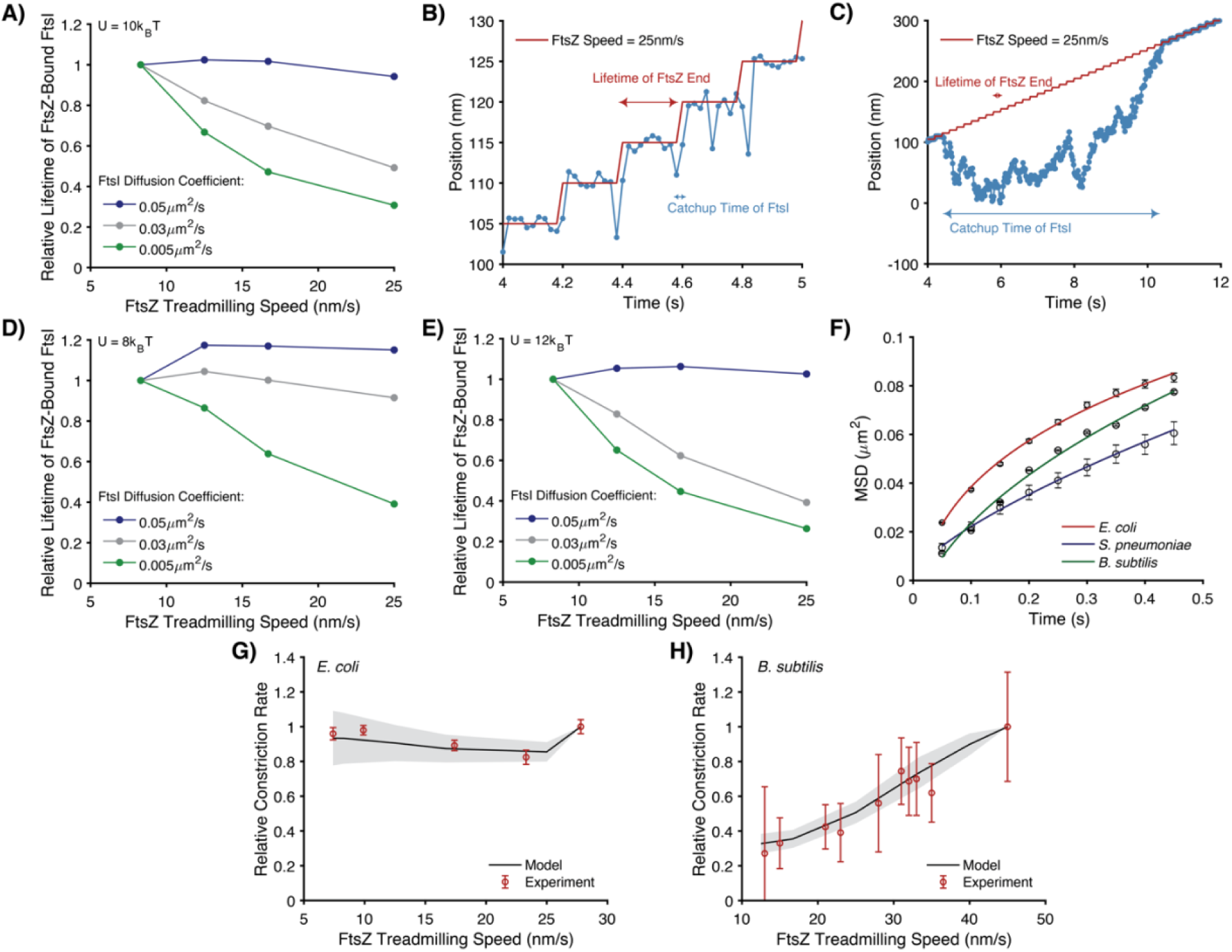
Dependence of sPG synthase’s processivity on FtsZ’s treadmilling speed is modulated by the synthase’s diffusion and binding potential. A) Predicted dependence of FtsZ-bound FtsI’s lifetime on FtsZ treadmilling speed with a binding potential of 10 *k*_B_T. B) A representative trajectory of FtsI’s persistent end-tracking when FtsI diffusion is fast (0.05 μm^2^/s). C) A representative trajectory of FtsI’s persistent end-tracking when the FtsI diffusion is slow (0.005 μm^2^/s). D) Predicted dependence of FtsZ-bound FtsI lifetime on FtsZ treadmilling speed with a weaker binding potential (8 *k*_B_T). E) Predicted dependence of FtsZ-bound FtsI lifetime on FtsZ treadmilling speed with a stronger binding potential (12 *k*_B_T). For A, D, and E, the initial condition is FtsI being randomly positioned along a FtsZ filament. The lifetime of a FtsZ-bound FtsI molecule was defined as the first time that the FtsI molecule escapes from the either end of the FtsZ filament for greater than 100 nm. For B and C, a fast diffusion allows FtsI to catch up with the shrinking end of FtsZ almost immediately, whereas it takes a long time for FtsI to catch up (if it eventually catches up) when it diffuses slowly. Here, the simulation time step is 10^−5^ s, the model results are plotted are plotted every 2×10^−2^ s. F) Measured mean-squared displacement (MSD) of FtsI, PBP2b and FtsW in wildtype *E*. *coli, B*. *subtilis*, and *S*. *pneumoniae* cells outside of the septal regions respectively. All three MSD curves were fitted using the function MSD = 4Dt_α_ since these molecules diffuse on a 2D curved membrane surface. From these data, the diffusion constants of FtsI, Pbp2b, and FtsW were extracted at 0.041, 0.038, and 0.028 µm^2^/s, with α = 0.29, 0.51, and 0.71, respectively. Error bars represent standard error of the mean. G) Model fitting of the dependence of sPG synthesis activity on FtsZ treadmilling speed in *E*. *coli*. The relative sPG synthesis activity was taken from previous constriction rate measurements in Coltharp et al^42^. and Yang et al^13^. H) Model fitting of the dependence of sPG synthesis activity on FtsZ treadmilling speed in *B*. *subtilis*. The relative sPG synthesis activity was taken from previous cell division time measurements in Bisson-Filho et al^12^. For G and H, the model fitting used the measured diffusion constants as that in F, varied the binding potentials, and determined their upper and lower limits (8 and 9 *k*_B_T for *E*. *coli*., and 10 and 12 *k*_B_T for *B*. *subtilis*), which mark the boundaries of the shaded regions.

The physical reason behind this drastic difference between fast and slow FtsI diffusion lies at the core of Brownian ratchet mechanism. Fast diffusion will allow FtsI to catch up with the shrinking end of an FtsZ filament in a very short time (Fig. 5B). When FtsI’s diffusion becomes slower and slower, it eventually becomes the rate-limiting factor in the Brownian ratchet – a slow FtsI molecule falls behind the FtsZ shrinking end and takes a long time to catch up with the departing FtsZ filament, or simply diffuses away and become lost (Fig. 5C). As such, further increasing the FtsZ treadmilling speed in the latter case will significantly reduce the chance of FtsI keeping up with the FtsZ shrinking end and, hence the lifetime of the FtsZ-bound FtsI. Crucially, this diffusion-modulated dependence of sPG synthesis on FtsZ treadmilling speed hinges on the binding between FtsI and FtsZ. When the binding potential is reduced, the lifetime of FtsZ-bound FtsI is less sensitive to FtsZ speed than its higher-potential counterpart (compare Figs. 5A, 5D and 5E). Therefore, how FtsZ’s treadmilling speed modulates the rate of sPG synthesis depends on the combined effects of the sPG synthase’s diffusion coefficient and FtsZ-binding potential.

To examine this hypothesis, we performed fast frame-rate single-molecule tracking to measure the diffusion coefficients of free sPG synthase molecules outside the septum (FtsI in *E*. *coli* and PBP2b in *B*. *subtilis*, Fig. 5F). As FtsZ only localizes to the midcell during cell division, sPG synthase molecules not localized to the midcell are considered free and not interacting with FtsZ. Using the measured diffusion coefficients, we then fit the experimentally observed dependence of cell wall constriction rate on FtsZ’s treadmilling speed in *E*. *coli* and *B*. *subtilis*^*12,42*^ with the normalized off-rate calculated from the model. The only free parameter in the model fitting is the binding potential, which was not possible to measure accurately in live cells with available experimental methods. As shown in Figs. 5G and 5H, we found that with the apparent diffusion constants of FtsI in *E*. *coli* and PBP2b in *B*. *subtilis* measured at ∼ 0.041 ± 0.0051 (mean ± S.E.M., N = 5049 trajectories) and 0.038 ± 0.0019 µm^2^/s (mean ± S.E.M., N = 6765 trajectories) respectively, and binding potentials set at 8-9 and 10-12 *k*_B_T, respectively, the model quantitatively recapitulated the differential dependence of cell wall constriction rate on FtsZ’s treadmilling speed in the two species as previously measured. The higher binding potential of PBP2b to FtsZ in *B*. *subtilis*, likely due to the significantly different protein-protein interactions in the Gram-positive bacteria, renders tighter coupling between end-tracking PBP2b molecules with FtsZ than that in *E*. *coli*, hence the fraction of end-tracking FtsI can be sensitively modulated by FtsZ’s treadmilling speed in *B*. *subtilis*, but that of FtsI in *E*. *coli*. cannot.

As a comparison, we also measured the diffusion of FtsW in *S*. *pneumoniae* (Fig. 5F). We found that the apparent diffusion coefficient of FtsW was at 0.028 ± 0.0004 µm^2^/s (mean ± S.E.M., N = 21 trajectories), in the same order of magnitude as that of *E*. *coli* and *B*. *subtilis*. As FtsW does not follow the treadmilling of FtsZ at all in *S*. *pneumoniae* at all, it is most likely that the binding potential between FtsW and FtsZ is significantly lower than 5 K_B_T as predicted by the model (Fig. 2A) under the experimental condition. It remains interesting to investigate in *S*. *pneumoniae* whether other divisome proteins are also independent of FtsZ’s treadmilling, or they exhibit conditional dependence once the protein-protein interactions of the divisome are altered due to the presence or depletion of their binding partners. These possibilities will be further investigated in our future work.

## Discussion

In this work, we presented data supporting a Brownian ratchet model that couples the directional movements of sPG synthases to FtsZ’s treadmilling and underlies the differential sensitivity of sPG synthesis to FtsZ’s treadmilling speed in *E*. *coli* and *B*. *subtilis*.

We first show that an sPG synthase molecule (here using FtsI as the model enzyme) can follow a treadmilling FtsZ polymer by end-tracking its shrinking tip but not the growing tip, due to the intricate interplay between the sPG synthase molecule’s diffusion and its binding potential to FtsZ. Only within a particular diffusion range (∼0.001 to 0.1 μm^2^/s) and at a sufficient binding potential (> 5 *k*_B_T), a sPG synthase molecule can exhibit FtsZ-treadmilling-dependent directional movement (Fig. 2). Furthermore, we show that the persistence run duration and distance of FtsI exhibit different dependence on FtsZ’s treadmilling speed, as predicated by the Brownian ratchet model (Fig. 3). Using single-molecule tracking, we confirmed these model predictions (Fig. 4). The ability of treadmilling FtsZ polymers to modulate the persistence run duration and distance of sPG synthase molecules could play an important role in regulating the spatial distribution of sPG synthases to ensure the correct septum shape. Finally, we show that the Brownian Ratchet model could explain the differential dependence of sPG synthesis activity on FtsZ’s treadmilling speed in *E*. *coli, B*. *subtilis* and *S*. *pneumoniae*.

Given experimentally measured diffusion coefficients of sPG synthases in different bacterial species, the Brownian ratchet model predicts that the tighter binding between PBP2b and FtsZ in *B*. *subtilis* could cause tighter coupling between them. Hence, the fraction of time a PBP2b molecule spends on FtsZ, and consequently the fraction of time it is off FtsZ to become available for sPG synthesis, can be sensitively modulated by FtsZ’s treadmilling speed. In *E*. *coli* or *S*. *pneumoniae*, the binding potential between FtsI/FtsW and FtsZ may not be as high as that in *B*. *subtilis*, and hence the fraction of time a FtsI/FtsW molecule remains end-tracking FtsZ exhibits much less sensitivity or no sensitivity at all to FtsZ’s treadmilling speed. Different binding potentials between these different bacterial species are likely, as detailed molecular interactions among the septal ring complexes are distinct in each species. These results suggest that the same Brownian-ratchet machinery may be at work but operate in distinct regimes of the parameter space in different species. Consequently, sPG synthesis depends on FtsZ treadmilling differentially, reflecting different strategies to meet different functional needs.

We note that additional factors could also be at play. For example, the expression level of sPG synthase relative to that of FtsZ in *B*. *subtilis* is significantly higher than that in *E*. *coli* or *S*. *pneumoniae*^43-51^ (Table S1). This suggests that the number of sPG synthases per FtsZ filament in *B*. *subtilis* would be higher than that in *E*. *coli* or *S*. *pneumoniae*. Our Brownian ratchet model predicts that this condition further enhances the sensitivity of sPG synthesis activity to FtsZ’s treadmilling speed, as sPG synthase molecules bound to the inner positions of FtsZ polymer would “knock” the end-tracking one off FtsZ (or vice versa, Fig. S1). Therefore, the faster the FtsZ treadmills, the faster the FtsZ shrinking end catches up to the sPG synthase in the middle and the more sPG synthase molecules will be dissociated from FtsZ to become available for sPG synthesis. This is a stark contrast to the case of a single sPG synthase per FtsZ filament, which is mostly likely the case in *E*. *coli* or *S*. *pneumoniae* (Fig. S1).

Moreover, the level of cell wall synthesis precursors, for example, could be another important factor. It is possible that across bacterial species, sPG synthase molecules are coupled to FtsZ’s treadmilling dynamics and their lifetime on FtsZ can be sensitively modulated by FtsZ’s treadmilling speed. However, if the level of a cell wall synthesis precursor is limiting, which is likely the case in *E*. *coli*, such a sensitivity could be further masked by the limited level of the precursor^52-55^. In *S*. *pneumoniae*, besides a low binding potential between the sPG synthase and FtsZ, cell wall synthesis precursor levels could also play a role in the independence of FtsZ’s treadmilling. High enough levels of PG precursors could saturate all sPG synthase molecules so that no free ones are available to track with FtsZ polymers.

In summary, given the lack of linear stepper motors in prokaryotic world, Brownian ratcheting appears to be an ancient mechanism for directed cargo transportation in bacteria – another salient example is ParA-mediated DNA partitioning^56-58^. Interestingly, a similar Brownian ratchet mechanism also underlies the directional movement of mitotic chromosomes by end-tracking spindle microtubule in eukaryotes^59^. Can we distill unified fundamental principle(s) by which evolution shapes the same Brownian ratchet mechanism to meet distinct needs under different contexts? We will relegate these exciting questions to our future study.

## Acknowledgements

The authors would like to thank Dr. S. Shaw and members in the Xiao and S. Holden labs for helpful discussions and technical assistance, Dr. G. Hauk for sharing plasmids and the CRISPR-Cas9/λ-red recombineering cloning method, R. McQuillen for help cloning pRM027, Dr. E. Goley for help with growth curve measurements, and Dr. L. Lavis for sharing JF646. This work was supported in part by NIH GM007445 (to J.W.M.), NIH R35 GM131767 (to M.E.W.), equipment grant NIH 1S10OD024988-01 (to Indiana University Light Microscopy Imaging Center), NIH F31AI138430 (to M.M.L.), NSF GRFP DGE1144152 (to G.R.S.), NIH R01 GM086447 (to J.X.), GM125656 (subcontract to J.X.), NSF EAGER Award MCB-1019000 (to J.X.), a Hamilton Smith Innovative Research Award (to J.X.), Johns Hopkins University Startup fund (to J.L.) and Catalyst Awards (to J.L.).

## Author Contributions

J.W.M., X.Y., G.R.S., K.E.B., M.M.L., and Z.L. performed experiments. J.L. carried out theoretical modeling. All authors contributed to concept development, data analysis, and manuscript writing.

