## Supplementary_Information for "Treadmilling FtsZ polymers drive the directional movement of sPG-synthesis enzymes *via* a Brownian ratchet mechanism"

### Section 1. Analytic solution of the dependence of FtsI's persistent run distance and duration on FtsZ's treadmilling speed.

We aim to obtain the analytical solution for the FtsZ treadmilling speed-dependence of run distance and duration of FtsI persistent end-tracking. A FtsI's persistent end-tracking trajectory can be decomposed into repeating steps, each of which consists of two consecutive processes. Process (1) is "stay-on": The FtsI molecule stays inside the binding potential of the FtsZ end subunit. Process (2) is "catch-up": The FtsI molecule catches up with the next FtsZ subunit in the row when the FtsZ end subunit dissociates. We next calculate the probabilities of the two processes.

For FtsI in the "stay-on" process (Fig. S1A), it can undergo two reactions in parallel: FtsI can escape from the binding potential of the FtsZ end subunit with the rate of $1/\tau_D$, and the FtsZ end subunit can dissociate with the rate of $1/\tau_Z$. Here, $\tau_Z$ is the average lifetime of FtsZ subunit, inversely proportional to the FtsZ treadmilling speed, $\tau_Z = \frac{5\ nm}{V_Z}$. $\tau_D$ is the average duration of FtsI staying in the binding potential if the FtsZ subunit never falls off. The probability that the FtsI is still in "stay-on" at time, *t*, is $P_1 = exp\left(-\left(\frac{1}{\tau_D}+\frac{1}{\tau_Z}\right)t\right)$. Thus, by the time the FtsZ end subunit dissociates, $P_1$ scales as $P_1 \sim \exp\left(-\frac{\tau_Z}{\tau_D}\right)$. Likewise, the catch-up probability of the FtsI molecule $P_2$ scales as $P_2 \sim \exp\left(-\frac{\tau_C}{\tau_Z}\right)$. Here, $\tau_C$ is the average catch-up time (Fig. S1B). Taken together, the probability for the FtsI end-tracking in each repeating step is:

$$P = P_1 P_2 \sim \exp\left(-\left(\frac{\tau_Z}{\tau_D}+\frac{\tau_C}{\tau_Z}\right)\right) \qquad \text{Eq. [S1]}$$

It follows that the probability of persistent end-tracking exact *N*-repeating steps is $P^N(1-P)$. Consequently, the average number of the FtsI persistent end-tracking steps is:

$$\langle N \rangle = \frac{\sum_{N=0}^{\infty} N \cdot P^N \cdot (1-P)}{\sum_{N=0}^{\infty} P^N \cdot (1-P)} \qquad \text{Eq. [S2]}$$

Taking the continuum limit, $\langle N \rangle = \frac{\int_0^\infty \{N \cdot P^N \cdot (1-P)\} dN}{\int_0^\infty \{P^N \cdot (1-P)\} dN}$, which yields $\langle N \rangle = \frac{\tau_D \tau_Z}{\tau_Z^2 + \tau_C \tau_D}$. The corresponding average run length and duration are:

$$\langle L \rangle = L_0 \frac{\tau_D \tau_Z}{\tau_Z^2 + \tau_C \tau_D} \qquad \text{Eq. [S3]}$$

and

$$\langle T \rangle = \frac{\tau_D \tau_Z}{\tau_Z^2 + \tau_C \tau_D} \tau_Z \qquad \text{Eq. [S4]}$$

Here, $L_0$ = 5 nm, the length of FtsZ subunit.

Specifically, the persistent end-tracking entails that $\tau_D >> \tau_Z >> \tau_C$. According to our numerical simulation, normally $\tau_D \geq 60\ s$ and $\tau_C \leq 0.001\ s$. Within this parameter range, the Eqs. [S3-4] can quantitatively recapitulate the distinctive dependences of run length and duration on FtsZ treadmilling speed. Herein Fig. S1C presents a representative result of the analytic solution with $\tau_D = 60\ s$ and $\tau_C = 0.0003\ s$.

Qualitatively speaking, the longer the lifetime of FtsZ subunit, FtsI in Process (1) will have a higher chance to escape the binding potential of FtsZ end subunit, decreasing the stay-on probability. In contrast, a longer lifetime of an FtsZ subunit will allow a longer time for the FtsI in Process (2) to catch up the next FtsZ subunit in the row, increasing the catch-up probability. Therefore, the balance between the stay-on and catch-up processes defines an optimal lifetime of an FtsZ subunit – and, hence, the optimal FtsZ treadmilling speed – that maximizes the probability of FtsI end-tracking per repeating step. This determines the maximum of the average number of repeating steps and explains the biphasic dependence of the FtsI persistent run length on FtsZ treadmilling speed. On the other hand, the duration per repeating step is approximately the lifetime of FtsZ subunit. As the FtsZ treadmilling speed increases, the decrease in the lifetime of FtsZ subunit outweighs the corresponding variation in the average number of repeating steps, leading to the decrease in the overall duration of persistent end-tracking.

### Section 2. Methods and Materials

#### Methods

**Media, Bacterial Strains, Plasmids**

Cells were grown in M9 minimal media or lysogeny broth (LB) (10% tryptone, 10% NaCl, 5% yeast extract). Fresh LB plates of strains were struck with appropriate antibiotics (detailed below) once-per-week from frozen stocks, and all cultures were started with a single colony. Bacterial strains and plasmids used are detailed in Table S1.

**Growth Curve**

Three biological replicates from TB28 and JM136 were grown from single colonies in M9+Glucose (M9) [1x M9 salts (Sigma-Aldrich M9956), 1x Amino Acids (Sigma-Aldrich M5550), 0.4% glucose, 1x Vitamins (Sigma-Aldrich M6895), 2mM $MgSO_4$, 100µM $CaCl_2$] as overnights at 30°C. The following day, the overnights' $OD_{600}$ were measured with a nanodrop and diluted to an $OD_{600}$ of 0.1 in 200µl M9 using a Corning Costar sterile 96 well plate. The 96-well plate was incubated in a Tecan Infinite M200 Pro set at 30°C, where it would measure the $OD_{600}$ of designated wells once every 30min for 23.5 hours, shaking the plate for 3min at 220rpm before measuring. To obtain the growth rate, the linear phase of the natural-log transformed growth curve data was fitted to a straight line. The slope of that line was used to calculate the doubling time through the equation below.

$$Doubling\ Time = \frac{\log_{10}(2)}{slope}$$

**Western blot of FtsI**

TB28 and JM136 were grown as overnights from single colonies in M9 at 37°C. They were propagated 1:50 in 3ml M9 and grown an additional 16hr at 25°C until their $OD_{600}$ reached ~0.5, at which point 500µl was harvested. These samples were pelleted, flash-frozen, and stored at -80°C for one hour. These pellets were then resuspended with PBS and mixed with 2x SDS buffer (100mM Tris-HCL, 4% SDS, 0.2% bromophenol blue, 20% glycerol, 100mM DTT). Cells were incubated for 10min at 95°C in a thermocycler, then 20µl of each were loaded into a 10% BioRad polyacrylamide gel with 5µl of PageRuler as a ladder. After electrophoresis, the gel was transferred at 25V for 2 hours to a nitrocellulose membrane. Membranes were checked for even transfer with a Ponceau stain before blotting. Rabbit α-FtsI, generously donated by Dr. David Weiss, was used as the primary antibody with a 1/50,000 dilution. Blots were then stained with 1/50,000 HRP goat anti-rabbit secondary antibody. Bands were visualized with a Clarity Western ECL Substrate BioRad kit and recorded using Blu-C autoradiography film for 30s. This western was repeated with three more biological replicates for each strain with the same result.

**Guide RNA and repair oligo design for making the Halo-FtsI$_{SW}$ fusion**

The chromosomal *ftsI* gene coordinates were first located using EcoCyc, which were then used to obtain the full *ftsI* gene sequence from NCBI (MG1655 genome, accession number NC_000913). This sequence was copied into ChopChop[1] to identify candidate protospacers near the N-terminus of FtsI covering residues 18 and 19 to make the sandwich fusion. The repair oligo was designed for λ-Red insertion[2] of HaloTag to break apart the protospacer, with 50nt overlap on either end of the codons for residues 18-19 (Table S3). Silent point mutations with comparable codon usage were also picked in this homology arm to edit the sequences of the protospacer and PAM to help select for a successful insertion of HaloTag.

**Plasmid Construction**

pJM44 was constructed through using InFusion Cloning. Briefly, a pACYC-sgRNA backbone was obtained from Dr. Glenn Hauk and inverse PCR was used to create a backbone (Table S3). A short oligo was designed to insert the 20bp protospacer in front of the guide RNA sequence, and an InFusion reaction was run to insert the sequence based on 15nt homology overlap with the template. The plasmid was then transformed into chemically competent Stellar competent cells (*E. coli* HST08) and outgrown in SOC for 1hr at 37°C. Cells were plated on LB + 150µg/ml chloramphenicol (CAM) and incubated overnight at 37°C. Colonies were screened by colony PCR of the sgRNA and confirmed via sanger sequencing. The plasmid was purified via ThermoScientific's GeneJET Plasmid Miniprep Kit.

**JM136 Construction**

CRISPR/Cas9 was performed with a method similar to previous work[3] TB28 harboring pKD46 and pJM25 was grown at 30°C overnight in LB + 60µg/ml carbenicillin (CB) + 50µg/ml kanamycin (KAN). Those cells were diluted the following day 1:100 in 50ml LB + 60µg/ml CB +

50µg/ml KAN and grown for 2hr or until their $OD_{600}$ reached ~0.5. Once the culture hit the proper $OD_{600}$, 0.2% arabinose was added to the culture to induce Cas9 for one hour. After induction, these cells were prepared to be electrocompetent such that fresh, concentrated *E. coli* could be immediately electroporated with 2µl concentrated sgRNA plasmid pJM44 and 10µl repair oligo. Cells were outgrown for 1.5hr at 30°C in SOC media then plated on LB + 50µg/ml CB + 50µg/ml KAN + 150µg/ml CAM to incubate for 2 days at 30°C.

Colonies were screened by colony PCR of the chromosomal locus of *ftsI*. Overnight cultures were started of any hits at 37°C in LB + 0.2% arabinose + 6% sucrose to kick out all plasmids. Subsequently, these overnights were serial-diluted to a factor of $10^{-6}$ and plated on LB plates that were then incubated overnight at 37°C. Twenty random colonies were selected and screened to ensure all plasmids were excised. A final colony with all three plasmids kicked out would be checked by PCR once more and sequenced to ensure that the locus was correct.

**pRM027 Construction**

The plasmid pRM027 ($P_{T5}$-*lac*::*meos3.2-ftsI*, *aph*) was constructed by amplifying *meos3.2*[4] and *ftsI* genes from pJB106 and pVS155-FtsI[5], respectively, and inserted to the linearized backbone from vector pCH027[5] using the Infusion protocol (Clontech Inc). The cat cassette was then replaced by an *aph* cassette amplified from pKD13.

**Preparing cells for single molecule tracking**

JM136 harboring pXY018 was grown overnight from a single colony at 37°C in M9 + 150µg/ml CAM. The following day, the culture was propagated 1:100 into fresh M9 +150µg/ml CAM and grown for 16hr at 25°C. The following morning, 3ml of culture was harvested at mid-log phase. Cells were concentrated to 100µl and incubated with 10nM of Janeliafluor 646 (JF646) for 30min. After labeling, JM136 was washed three times with M9 medium without vitamins ($M9_{-}$) and concentrated to 50µl.

EC812 harboring pRM027 was grown overnight from a single colony at 37°C in LB + 150µg/ml CAM and 50µg/ml KAN with 0.2% L-arabinose. The culture was reinoculated 1:100 into fresh M9 media overnight at room temperature to log-phase for imaging. mEos3.2-FtsI supports cell growth and single molecule tracking at basal-level expression.

A 3% agar pad was prepared using a nanopillar chip with pillars of a diameter range of 1.2-1.4μm and a length of 4.5μm. After cooling for 30min, cells were added to the agar pad and incubated for 2min. The agar pad was then washed by adding 1ml M9 and incubating the agar pad for 2min. The media was aspirated off and the agar pad set out to dry at room temperature for ~20min. The agar pad was sandwiched with a coverslip, sealed in a Bioptechs FSC2 chamber, and taken to the microscope for imaging.

**Microscope and imaging setup**

JM136 harboring pXY018 was imaged using two split channels in widefield on an Olympus IX-71microscope. JF646 was imaged with 647 set to ~50W/cm$^2$ and GFP-ZapA was imaged with 488 set to ~5W/cm$^2$. The channels were split with an Optosplit II system containing 600rdc, with a 700/55 emission filter for JF646 and 540/30 emission filter for GFP-ZapA. We used an Andor iXon 897 Ultra EM-CCD camera with an APON100xOTIRF objective (1.49NA/oil) and engaged 1.6x optivar. Our camera's EM-Gain was turned on to 300 with a pre-amplifier gain setting on 3 and digitizer set to 16-bit. Baseline clamp was activated, with baseline offset set to 100.

**Phase contrast imaging**

TB28 and JM136 were grown overnight from single colonies at 37°C in M9. The following day, the cultures were propagated 1:100 into fresh M9 and grown for 16hr at 25°C until mid-log phase. 500μl of each culture was pelleted and concentrated in 50μl. 0.5μl of each strain was added to separate 3% M9 agar pads (pre-set for 30min before use), sealed in a BiopTechs FSC2 chamber, and taken to the microscope for imaging. Cells were imaged with phase contrast on an Olympus IX-71 microscope with a 100x/1.30NA Oil Ph3 objective and engaged 1.6x optivar. Images were recorded at a full 512x512 region on an Andor iXon Ultra Em-CCD camera (above) at 100nm/pixel. Phase contrast images were recorded with 100ms exposures and processed with Oufti to measure cell length[6].

**Single molecule tracking of Halo-FtsI::JF646**

All microscopy was recorded using Metamorph software. A region of 300 pixels wide and 512 pixels tall was used to capture both channels. Samples were put on the microscope and acclimated for 30-60min before imaging began to minimize axial drifting. 25 regions were selected with cells in microholes, and an automated journal would run to 1) autofocus on cells lying on the surface of the agar pad, then 2) move up into the sample 2μm to find the Z-ring of cells in microholes and 3) autofocus on the GFP-ZapA ring. Once a focal plane was set, the journal would record a 400-frame movie, where each frame encapsulated 1 second (500ms exposure, 500ms dark time).

#### Aligning channels to correct for chromatic aberration

To correct for chromatic aberration, we imaged TetraSpeck fluorescent beads in both channels using a 50ms exposure with the same 300x512 region. 647 remained set to ~50W/cm$^2$. 488 was set to ~40W/cm$^2$. To align the channels, we cropped the two using a custom Matlab script that then used the imregister function to align the 488 (ZapA-GFP) channel to the 647 channel (JF646). The dimensions of the crop and the transformation matrix from this alignment were used to crop all channels and align the GFP-ZapA channel to JF646.

#### Analyzing single molecule trajectories

Cropped 647 channels were first processed with ThunderSTORM[7], a plug-in for ImageJ[8]. Image filtering used a wavelet filter (B-spline) with an order of 3 and scale of 2.0. A local maximum was used to localize the molecules with 1.5*std(Wave.F1) used to identify peak intensity threshold and a connectivity of 8-neigborhood. Sub-pixel localization used a guassian PSF with a 3-pixel fitting radius and an initial sigma of 1 pixel. The post-processed data was filtered to exclude intensity values less than 300 and a sigma bandpass filter of 60-300nm. All analysis thereafter used custom scripts in Matlab R2019a. The localizations were linked to trajectories using a nearest-neighbor algorithm modified from Sbalzarini and Koumoutsakos[9]. To link molecules which may have blinked across frames or left the focal plane, a time threshold of 15 frames was applied. The distance threshold was 300nm/1frame, which approximates to a diffusion coefficient of ~0.05μm$^2$/s, or a max speed of 300nm/s. Only trajectories with a corresponding GFP-ZapA ring were chosen for the next step.

GFP-ZapA stacks served as both a marker to autofocus for imaging as well as for estimating the Z-ring diameter. Maximum-intensity projections were taken of movies with cells expressing GFP-ZapA and then subsequently aligned to the 647 channel as described above. Cells of interest were cropped out and a circle was then fit to the intensity profile of the GFP signal. Using the diameter, the "real" position of FtsI along the cell envelope can be back calculated and estimated. The trajectories were then manually segmented into mobile states only when segments were a minimum of 4 frames (4 seconds) long with processive displacements in one direction. The selected segments were fit to a straight line to minimize noise and classified as “processive” or “stationary” based on the classification description detailed in the following section.

Processive segments were used to obtain the velocity, dwell time, and persistent length measurements.

#### Segmentation Classification

The trajectories were first segmented manually (a segment contains at least 4 data points and with consistent noise). After segmenting trajectories (Fig. S5), we classify whether the segments move processively or remain stationary. Included with our segmentation are a set of observables, $\{v, d, l, r\}$, where $v$ is the slope of a linear fit of the segment, $d$ is the total displacement, $l$ is the trajectory length, and $r$ is the standard deviation of all positions against the linear fit. Note these four parameters are not independent since $d = l \cdot v$. We subsequently combine the parameters to $\{v,\ l,\ R\}$ where, $R = \frac{r}{d}$, a dimensionless quantity representing the relative level of the residual to displacement (a noise to signal parameter). Through manual inspection, we determined to classify segments as processive based on a threshold of $R \leq 0.4$.

On average, one nanopillar experiment in a day (a biological replicate) will yield 0-5 trajectories. Single cells can give up to 1-3 usable trajectories for analysis. The 77 total trajectories used in this paper come from 18 biological replicates and 49 total cells. They yielded 232 total segments, 139 of which were marked as processive by this $R$ threshold.

#### Deconvolution of the FtsI Fast Population

The data presented in Fig. 4FGH were representative of the fast, inactive population of FtsI from a two-population fit to a log-normal CDF (Fig. S5BCD)[10].

$$f(x) = P\left(\frac{1}{x\sigma_1\sqrt{2\pi}}\right)e^{-\frac{1}{2}\left(\frac{\ln(x)-\mu_1}{2\sigma_1^2}\right)^2} + (1-P)\left(\frac{1}{x\sigma_2\sqrt{2\pi}}\right)e^{-\frac{1}{2}\left(\frac{\ln(x)-\mu_2}{2\sigma_2^2}\right)^2}$$

Where P is the percent makeup of the respective population. Briefly, we bootstrapped the FtsI data 200 times, iteratively fitting the CDF to obtain estimates for each fit parameter. A global fit of the mean bootstrapped values is presented in Fig. S5B, and individual fits for each population are shown with respect to FtsZ (from Yang et al.[11]) in Fig. S5C. We then used these parameters to overlay the raw FtsI histogram of 139 directional segments (Fig. S5D). Using the slow population fit, we resampled the FtsI data 100 times by removing segments that would be selected for a bin determined by the slow population's percent makeup compared to the original histogram.

The final histogram and scatterplots in Fig. 4FGH are representative plots from the resampling, where 74 segments remain that are likely to be fast-moving.

**Three-dimensional (3D) PALM single-molecule tracking and analysis**

Astigmatism-based 3D single-molecule tracking was performed on the same microscope. The 568nm excitation laser power was set to 500W/cm$^2$ with a 30ms exposure time for 5,000-10,000 frames of continuous acquisition. During the imaging, 0-1W/cm$^2$ 405nm activation light was increased stepwise and applied to activate mEos3.2-FtsI molecules. The UV power was tuned from sample to sample to maintain a low enough number of red-emitting molecules in each cell (<1 spot/frame/cell) for single-molecule localization and tracking.

Localization of single molecules was determined using the ThunderSTORM plugin in ImageJ[7,8]. Molecules with anomalous brightness or uncertainty (>3σ) were filtered out. Molecules were tracked across frames using custom Matlab scripts implementing the tracking algorithm described in Sbalzarini et al[9]. To calculate the MSD, all trajectories longer than 4 frames were selected, and the squared displacements were calculated in 2D. Consecutive frames in the trajectories were used for displacement calculation. The MSD curve was fitted by the anomalous diffusion equation: $MSD_{2D} = 4Dt^{\alpha} + D_0$, where $D_0$ reflects the localization uncertainty under the imaging conditions.

**Single Molecule Tracking of PBP2b in *B. subtilis***

*B. subtilis* strain bGS28, in which the native Pbp2B has been replaced with an IPTG-inducible, Halo-tagged Pbp2B, was imaged as described in Bisson-Filho et al[12]. Briefly, cells were grown in CH media, Pbp2B was induced with 20 μM IPTG and labeled with 100 nM JF549 conjugated to HaloTag ligand, and cells were immobilized under an agarose pad for imaging. Cells were imaged in TIRF on a Nikon N-STORM microscope; time lapses were acquired with streaming 30 ms exposures for 1 minute. Particle tracking was performed in TrackMate using the simple LAP tracker with the following settings: particle diameter was 300 nm, maximum linking distance was 300 nm, and no frame gaps were allowed. Tracks between 5 and 25 frames were analyzed further using custom MATLAB code[12]. MSD vs t was calculated for each track, and the diffusion coefficient was computed by least- squares fitting to $MSD_{2D} = 4Dt^{\alpha} + D_0$, where $D_0$ reflects the localization uncertainty under the imaging conditions.

### Single Molecule Tracking of FtsW in *S. pneumoniae*

S. pneumoniae strain IU15096 (D39 Δ*cps rpsL1 ftsW*-$L_0$-*ht*-$P_C$-*erm*) expresses a derivative of FtsW from its native chromosomal locus, with FtsW fused to a 10-amino acid linker ($L_0$) connected to the HaloTag (HT) domain[13]. Growth of cells labeled with JaneliaFluor 549 HaloTag ligand on agarose pads containing C+Y, pH 7.1 medium, and live cell imaging by TIRFm were performed as described previously[13]. Images were collected every 50 ms (20 FPS) for 100 to 200 s. The 561 nm (FtsW-HT) channel used a 21ms exposure, 100% T, and a TIRF angle of 91.2°, while the DIC channel used a 3 ms exposure, 50% T, and an angle of 0° (epi). Single cells were cropped from an image, and the FtsW-HT channel was processed using the FIJI plug-in ThunderSTORM[7,8]. Image filtering used a Wavelet filter (B-Spline) with an order of 3 and a scale of 2. The localization method was Local maximum, with a Peak intensity threshold of 1.5*std(Wave.F1) and 8-neighbourhood connectivity. Sub-pixel localization used a PSF: Gaussian method with a fitting radius of 3 pixels, fitting method of Weighted Least Squares, and an Initial sigma of 1 pixel.

Images depicting single molecule localizations of FtsW-HT from a single cell were visually inspected to ensure each frame had a maximum of one localization. Additional localizations resulting from single pixel noise were manually removed. A maximum of 5 consecutive frames without a localization was allowed within a single trajectory. From individual cells, time averaged mean square displacements (TA-MSD) were calculated on trajectories longer than 15 frames. The first third of each trajectory was used for further analysis.

Using TA-MSDs from individual cells, the diffusion coefficient (D) was calculated in two ways, giving similar results. (1) D was calculated for each lag time (t) using: $D_t$ = TA-MSD(t) / (4*t). $D_t$ was averaged over all lag times to generate D for each cell ($D_{cell}$), which were then averaged over all cells to give $D_1$. (2) In a plot of MSD versus lag time for each individual cell, the slope of linear regions was determined. The slope was divided by 4 to generate D for each cell ($D_{cell2}$), which were then averaged over all cells to give $D_2$. $D_1$ and $D_2$ are both reported as "Measurements from TA-MSD" (mean ± SD). Additionally, TA-MSDs for each lag time were averaged across cells, generating ensemble-averaged MSDs (EA-MSDs). EA-MSDs from over 20 cells (lag times between 0.05 and 0.55 s) were fit to a power series equation (GraphPad Prism): EA-MSD(t) = 4D*$t_\alpha$, to determine α = 0.69. $D_{t3}$ was calculated using: $D_{t3}$ = EA-MSD(t) / (4*$t_\alpha$), which was then averaged over all lag times and reported as the "Measurement from EA-MSD" (mean ± $S_{y.x}$, where $S_{y.x}$ is a goodness-of-fit descriptor similar to root mean square error).

### Section 3.  Supplemental Tables

| | Source | Abundance (relative value to FtsZ in parentheses) | | |
|---|---|---|---|---|
| | | FtsZ | FtsI/PBP2b/PBP2x | FtsW |
| *E. coli* | Arike et al.[14] | 585 (1) | 0.21 (0.00359) | 0.14 (0.00239) |
| | Krug et al.[15] | 1291 (1) | 1.97 (0.00152) | 2.12 (0.00164) |
| | PRIDE PRD000418[16] | 507 (1) | 13.8 (0.02722) | 1.97 (0.00389) |
| | Valgepea et al.[17] | 956 (1) | 36.3 (0.03797) | 27.4 (0.02866) |
| | Arike et al.[14] | 942 (1) | 59.0 (0.06263) | 12.6 (0.01337) |
| | Lu et al.[18] | 1376 (1) | 11.5 (0.00836) | 0.91 (0.00066) |
| | Lewis et al.[19] | 887 (1) | 3.83 (0.00432) | 0.86 (0.00097) |
| | PRIDE PRD000485[20] | 811 (1) | 27.5 (0.03391) | 6.01 (0.00741) |
| | PRIDE PRD000485[20] | 721 (1) | NA | 6.11 (0.00847) |
| | PAX db 4.1[21] | 833 (1) | 0.25 (0.00030) | NA |
| *B. subtilis* | Chi et al.[22] | 744 (1) | 87.5 (0.11761) | 53.0 (0.07124) |
| | PAX db 4.1[21] | 894 (1) | 97.5 (0.10906) | 19.9 (0.02225) |
| | PAX db 4.1[21] | 909 (1) | 98.5 (0.10836) | 16.6 (0.018262) |
| *S. pneumoniae* | Noirclerc-Savoye et al.[23] | NA | 260 | NA |
| | Lara et al.[24] | 3000 | NA | NA |
| | *E. coli* average relative abundance | | 0.01962 | 0.00726 |
| | *B. subtilis* average relative abundance | | 0.11168 | 0.03725 |
| | *S. pneumoniae* relative abundance | | 0.087 | NA |

**Table S1**. Comparison of PG synthase abundance between *E. coli*, *B. subtilis*, and *S. pneumoniae*. *E. coli* and *B. subtilis* data were gathered from the protein abundance database[21]. For *E. coli*, we pulled all results from searching "FtsI," "FtsW," or "FtsZ." For *B. subtilis* we pulled all results from searching "PBPb", "FtsW," or "FtsZ." *S. pneumoniae* estimates were obtained from the literature[23,24].

| **Strain** | **Genotype** | **Reference/Source** |
|---|---|---|
| TB28 | *E. coli* MG1655 *lacIYZ <> frt* | Bernhardt 2003[25] |
| JM136 | TB28 ftsI::halotag-$ftsI_{18-19}$ | This Study |
| EC182 | *E. coli* MG1655 *ftsI*::*cat* (*attL-lom*)::*bla lacIQ pBad-ftsI* | Wissel & Weiss 2004[26] |
| bGS28 | *B. subtilis* PY79 *pbp2B::erm-$P_{hyperspank}$-HaloTag-15aa-pbp2B* | Bisson-Filho 2017[12] |
| IU15096 | *S. pneumoniae* D39 Δ*cps rpsL1 ftsW*-$L_0$-*ht*-$P_C$-*erm* | Perez 2019[13] |

| **Plasmid** | **Genotype** | **Reference/Source** |
|---|---|---|
| pXY018 | pCA24N, *lacIQ*, *GFP-ZapA* | Yang 2017[11] |
| pRM027 | $P_{t5-lac}$::*meos3.2-ftsI* | This Study |
| pKD46 | repA101(ts), ParaB$_{-}$ (gam bet exo) | Datsenki 2000[2] |
| pGH33 | pET28a, $P_{bad}$::*spCas9*, *ampR* | This Study |
| pGH34 | pACYC, *cmR*, $P_{bad}$::*sacB*, $P_{bad}$::*sgRNA* | This Study |
| pJM44 | pACYC, *cmR*, $P_{bad}$::*sacB*, $P_{bad}$::*$ftsI_{18-19}$sgRNA* | This Study |

**Table S2**. Strains and plasmids used in this study.

| **Name** | **Purpose** | **Sequence** |
|---|---|---|
| JM26 | Amplifying pACYC vector (R) | GTGCACATTATACGAGCCGATGA |
| JM35 | Amplifying sgRNA insert (R) | CGGCGTAGAGGATCCACAGGACGGGTGTGGTCGC |
| JM44 | Amplifying pACYC vector (F) | ATCATGGCGACCACACCCGTCCTGTGGATCCTCTAC |
| JM97 | Amplifying *$ftsI_{18-19}$* sgRNA (F) | TCGTATAATGTGCACAACATGCCAACTTTATCAGTGTTTTAGAGCTAGAAATAGCAAGTTAA |
| JM95 | Amplifying *halotag-$ftsI_{18-19}$* repair oligo (F) | AAGCAGCGGCGAAAACGCAGAAACCAAAACGTCAGGAAGAACACGCAAACGGTGGCGGTGGTAGCGGATCCGAAATCGGTACTGGC |
| JM96 | Amplifying *halotag-$ftsI_{18-19}$* repair oligo (R) | AGCGCCAGGAGAATACAGCCGCATAACAACGCAAAACGCCATGAAATGAAGCTACCACCGCCACCGGAAATCTCCAGAGTAGAC |

**Table S3**. Primers used in this study. JM97 contains a 20nt protospacer for amplifying the sgRNA. JM95 and JM96 have GGGGS linkers encoded in their sequence in addition to the 50nt overhangs with the genomic site for λ-Red insertion.

| Position | Mutation | Annotation | Gene | Description |
|---|---|---|---|---|
| 257,908 | Δ776bp | | *insB9–[crl]* | *insB9, insA9, [crl]* |
| 361,267 | Δ6303bp | | *[lacA]–lacI* | *[lacA],lacY,lacZ,lacI* |
| 1,978,503 | Δ776bp | | *insB-5-insA-5* | *insB-5, insA-5* |
| 1,995,620 | IS*5* (–) +4b | Intergenic (-261/-32) | *uvrY* ← *I* → yecU | DNA-binding transcriptional activator UvrY/protein YecU |
| 2,173,363 | Δ2 bp | Pseudogene (915-916/1358nt) | *gatC* ← | Galactitol-specific PTS enzyme IIC component |
| 2,958,626 | C → T | A87T (GCA → ACA) | *ptrA* ← | protease 3 |
| 3,560,455 | +G | Pseudogene (151/758nt) | *glpR* ← | DNA-binding transcriptional repressor GlpR |
| 4,296,381 | +GC | Intergenic (+587/+55) | *glltP* → *I* ← *yjcO* | Glutamate/aspartate: H(+) symporter GltP/Sel1 repeat-containing protein YjcO |

**Table S4.** Identified mutations and deletions in the parent strain TB28 from whole genome sequencing in comparison to *E. coli* strain MG1655. JM136 shared these same perturbations in addition to insertion of HaloTag at FtsI's N-terminus.

### Section 4. Supplemental Figures

#### Figure captions

**Fig. S1**. Abundance of sPG synthase modulates FtsZ treadmilling speed-dependence of cell wall synthesis. A) Representative simulation trajectory of two sPG synthases on a treadmilling FtsZ filament. *Left*: Schematic of two sPG synthases on the same treadmilling FtsZ filament, as compared to the case of one sPG synthase. Note that as the two sPG synthases move along the one-dimensional track of a FtsZ filament, one of them will start to end-track the FtsZ shrinking end, with the hops around in the middle of the FtsZ filament. As the first pass, the only interaction between the sPG synthases considered by the current model is a short-ranged volume exclusion preventing them from crossing each other – when the two complexes touch each other, they "bounce" apart by 5 nm. As such, when the two sPG synthases approach each other, the end-tracking one will be "knocked" off the track and the other will take over the baton and start end-tracking. B) Dependence of FtsZ-bound sPG synthase lifetime on FtsZ treadmilling speed. C) sPG synthase levels modulate the dependence of sPG synthase activation rate (i.e., the inverse of the lifetime in (B)) on FtsZ treadmilling speed. D) sPG synthase level and FtsZ-binding potential govern the sensitivity of sPG synthase activation on FtsZ treadmilling speed. With the FtsZ treadmilling speed increasing 3-fold (from 8.3 to 25 nm/s), we counted it as "insensitive-dependence", if the corresponding changes in sPG synthase activation rate is less than 30%, relative to the activation rate at FtsZ treadmilling speed of 8.3 nm/s. Otherwise, we counted it as "sensitive-dependence". For (A – D), 100 independent stochastic simulation trajectories were simulated and averaged for each data point, wherever it applies. If not otherwise mentioned, the nominal parameter set is that the diffusion constant of free sPG synthases is 0.04 $\mu m^2/s$, sPG synthase-FtsZ binding potential 10 $k_B T$, and FtsZ treadmilling speed 25 nm/s.

**Fig. S2**. Model results of FtsI movement at the FtsZ growing end. A) A representative simulated trajectory of a slow-diffusing FtsI. It gets stuck at the FtsZ growing end until it can persistently end-track the FtsZ shrinking end. B) A representative simulated trajectory of a fast-diffusing FtsI.

It escapes from the FtsZ growing end before the FtsZ shrinking end approaches it. The diffusion constant of free FtsI is set to be 0.01 mm$^2$/s in A) and 0.04 mm$^2$/s in B). The rest of the parameters are the same for A) and B): FtsI-FtsZ binding potential is 10 kT and FtsZ treadmilling speed is 25 nm/s.

**Fig. S3**. Model results of analytic solution. A) Schematic of chemical reactions for the FtsI bound to the FtsZ end subunit. B) Schematic of chemical reactions for the free FtsI to catch up with the FtsZ end subunit. C) FtsZ treadmilling speed-dependence of the average run distance and duration of FtsI persistent end-tracking. Here, the plot is calculated based on the analytic solutions of Eq. S[3-4], where $\tau_D = 60\ s$ and $\tau_C = 0.0003\ s$. As long as $\tau_D >> \tau_Z >> \tau_C$, the essence of the plot holds up.

**Fig. S4**. A) Growth curve of TB28 vs. JM136 (with Halo-FtsI) in M9 minimal media at 30°C. Shaded region is the standard error. B) TB28 vs. JM136 doubling time, calculated from the growth curve in A. Error bars are standard deviation. C) Phase contrast images of TB28 and JM136. Scale bar is 5µm. D) Cell length of TB28 vs. JM136, measured from Oufti[6]. Error bars are standard deviation. E) Representative western blot of FtsI in TB28 and JM136. "L" denotes the lane the ladder was run in. F) Halo-FtsI localizes mid-cell in Halo-FtsI. Cells were stained with 1µM JF646. Scale bar is 5µm. (A,B,C and E) were performed with at least three biological replicates.

**Fig. S5**. Segmentation and deconvolution of FtsI directional segments. A) To determine the noise-to-signal ratio $R = \frac{r}{d}$, we use the residual and displacement information from fitted segments. B) A two-population CDF fit to the raw velocity data. Parameters were measured on the ln-scale, where P = 0.49 ± 0.15, $\mu_1$ = 2.1 ± 0.14, $\sigma_1$ = 0.50 ± 0.15, $\mu_2$ = 3.3 ± 0.20, $\sigma_2$ = 0.50 ± 0.12 (all mean ± S.E.M.). C) Fitting the two FtsI populations to their separate fits, with FtsZ from Yang et al.[11] for comparison. D) Raw FtsI histogram with the two-population overlay, used for deconvolution of the fast population.

**Fig. S1**

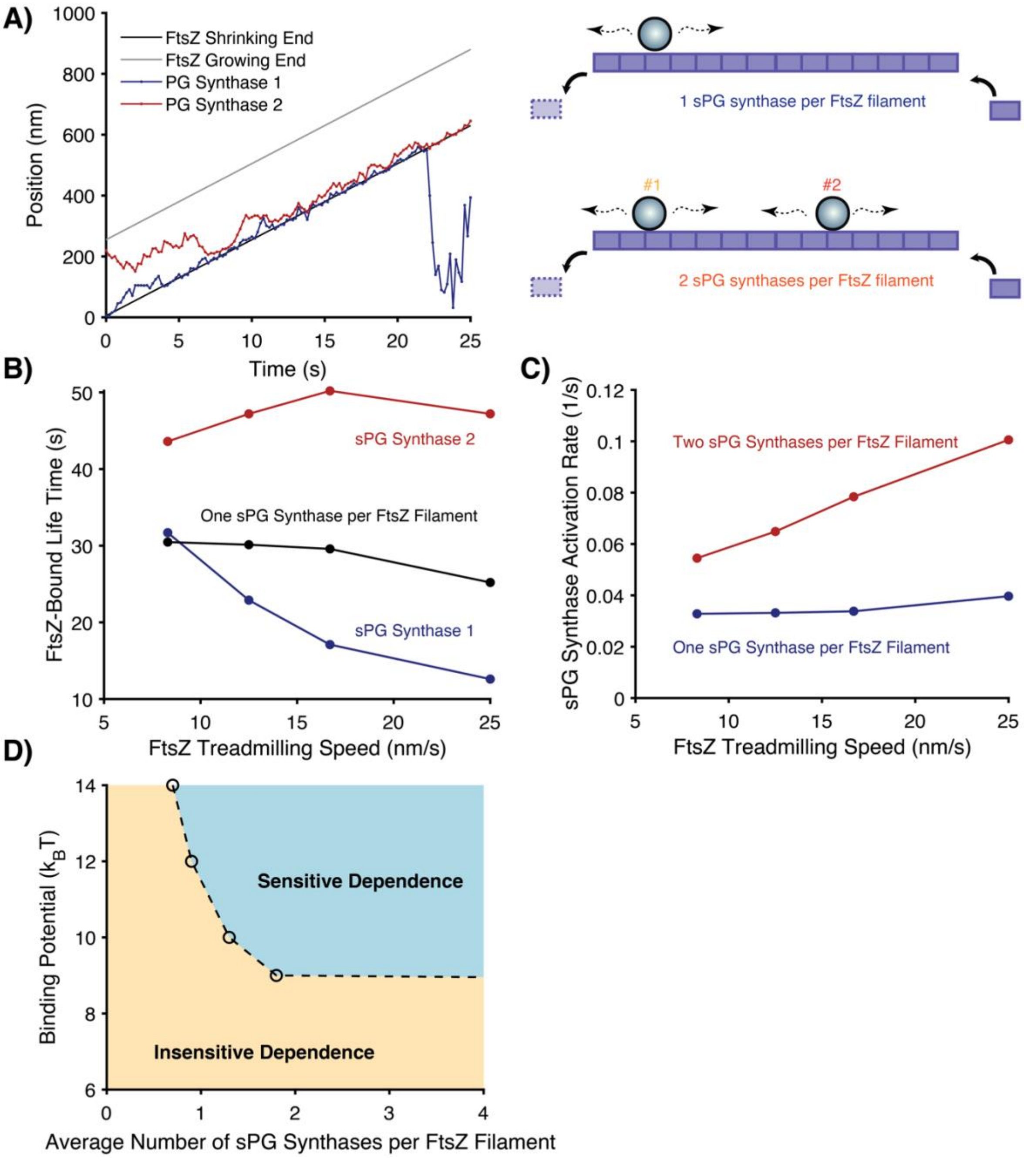
A)
FtsZ Shrinking End
FtsZ Growing End
PG Synthase 1
PG Synthase 2
Position (nm)
Time (s)
1 sPG synthase per FtsZ filament
#1
#2
2 sPG synthases per FtsZ filament
B)
FtsZ-Bound Life Time (s)
sPG Synthase 2
One sPG Synthase per FtsZ Filament
sPG Synthase 1
FtsZ Treadmilling Speed (nm/s)
C)
sPG Synthase Activation Rate (1/s)
Two sPG Synthases per FtsZ Filament
One sPG Synthase per FtsZ Filament
FtsZ Treadmilling Speed (nm/s)
D)
Binding Potential ($k_BT$)
Sensitive Dependence
Insensitive Dependence
Average Number of sPG Synthases per FtsZ Filament

**Fig. S2**

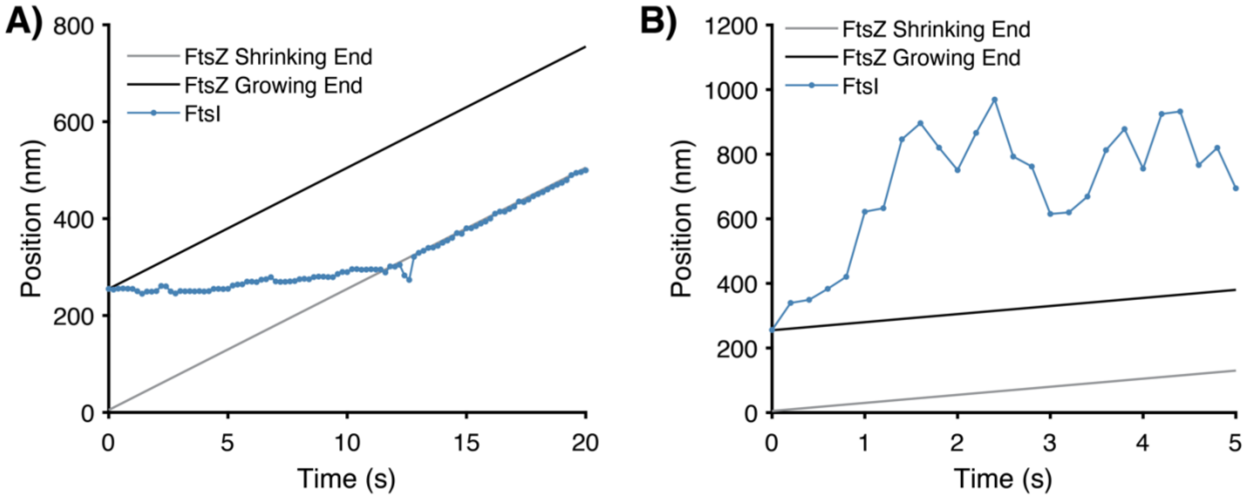
A)
800
600
400
200
0
Position (nm)
FtsZ Shrinking End
FtsZ Growing End
FtsI
0
5
10
15
20
Time (s)
B)
1200
1000
800
600
400
200
0
Position (nm)
FtsZ Shrinking End
FtsZ Growing End
FtsI
0
1
2
3
4
5
Time (s)

**Fig. S3**

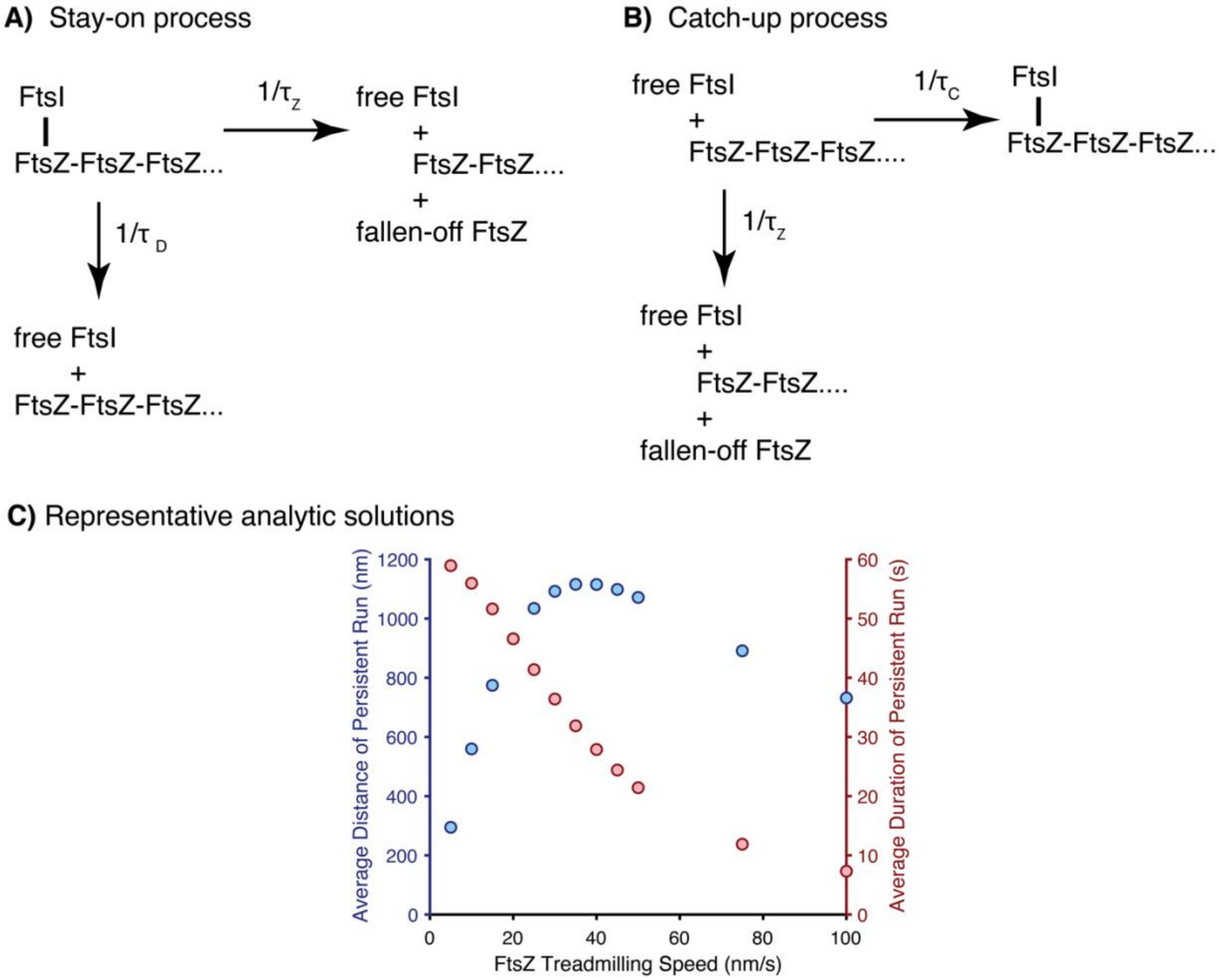
A) Stay-on process
FtsI
FtsZ-FtsZ-FtsZ...
1/τZ
free FtsI
+
FtsZ-FtsZ....
+
fallen-off FtsZ
1/τ D
free FtsI
+
FtsZ-FtsZ-FtsZ...
B) Catch-up process
free FtsI
+
FtsZ-FtsZ-FtsZ....
1/τC
FtsI
FtsZ-FtsZ-FtsZ...
1/τZ
free FtsI
+
FtsZ-FtsZ....
+
fallen-off FtsZ
C) Representative analytic solutions
Average Distance of Persistent Run (nm)
Average Duration of Persistent Run (s)
FtsZ Treadmilling Speed (nm/s)
0
200
400
600
800
1000
1200
10
20
30
40
50
60
0
20
40
60
80
100

**Fig. S4**

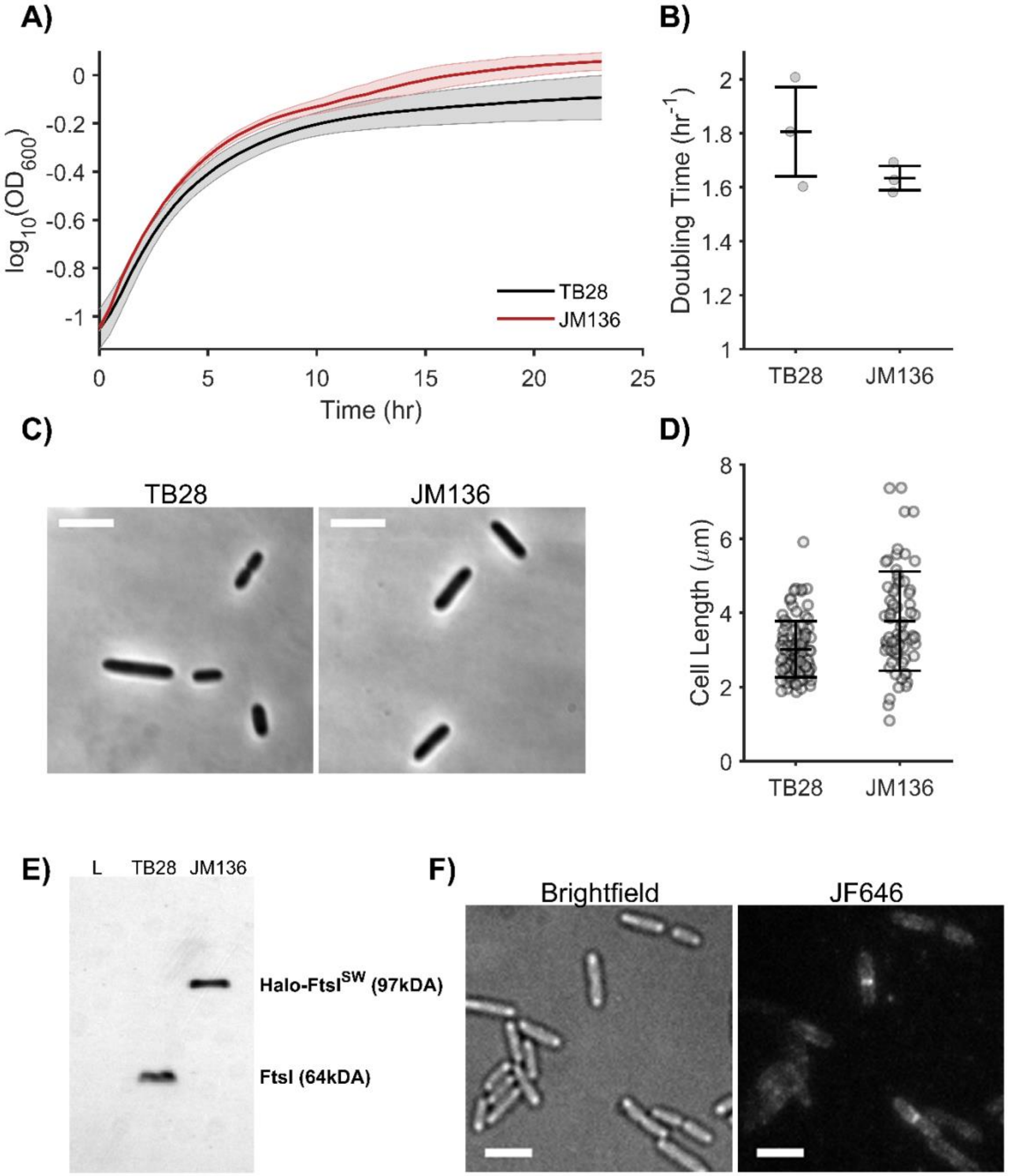
A)
0
-0.2
-0.4
-0.6
-0.8
-1
log10(OD600)
0
5
10
15
20
25
Time (hr)
TB28
JM136
B)
Doubling Time (hr-1)
2
1.8
1.6
1.4
1.2
1
TB28
JM136
C)
TB28
JM136
D)
Cell Length (μm)
8
6
4
2
0
TB28
JM136
E)
L
TB28
JM136
Halo-FtsISW (97kDA)
FtsI (64kDA)
F)
Brightfield
JF646

**Fig. S5**

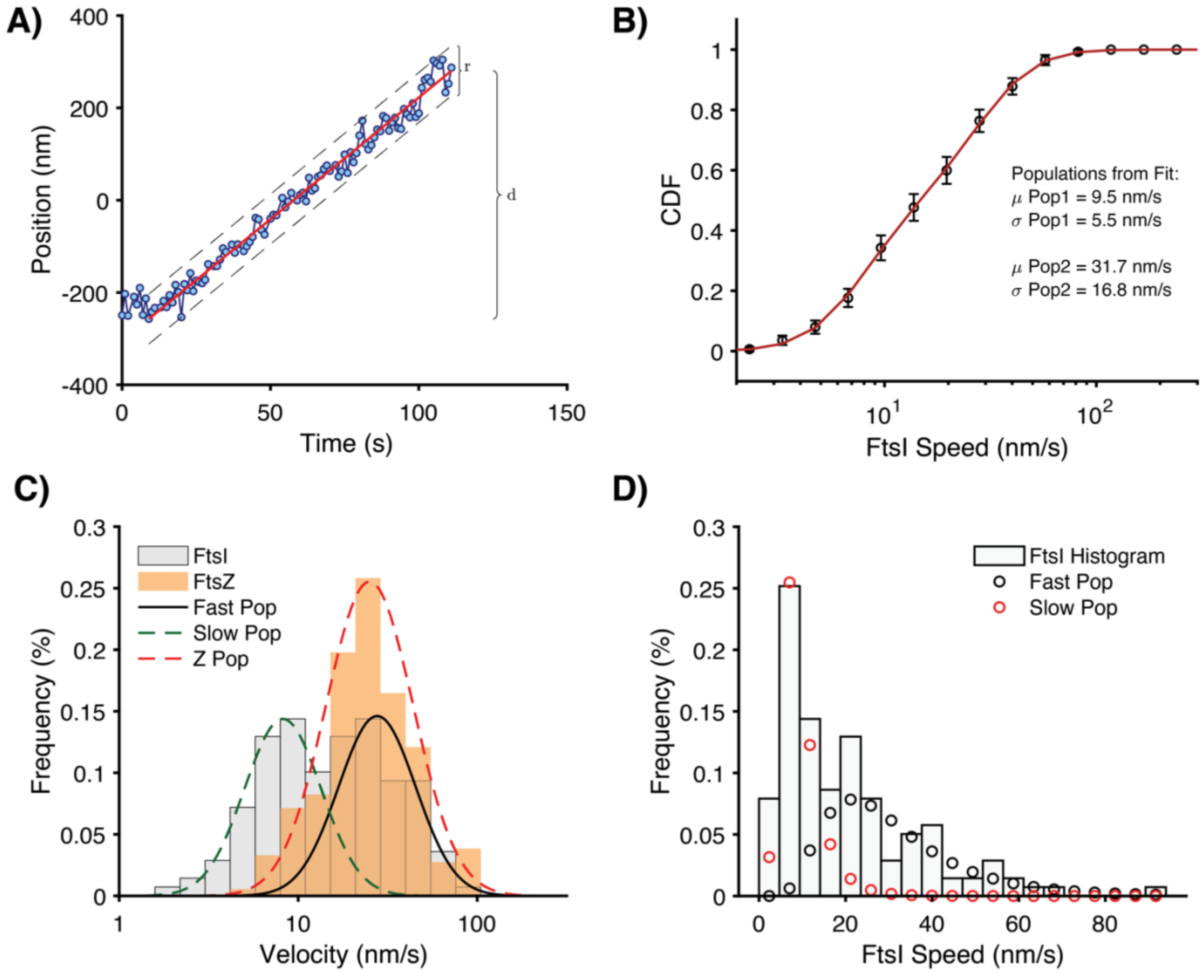
A)
Position (nm)
Time (s)
r
d
B)
CDF
FtsI Speed (nm/s)
Populations from Fit:
μ Pop1 = 9.5 nm/s
σ Pop1 = 5.5 nm/s
μ Pop2 = 31.7 nm/s
σ Pop2 = 16.8 nm/s
C)
Frequency (%)
Velocity (nm/s)
FtsI
FtsZ
Fast Pop
Slow Pop
Z Pop
D)
Frequency (%)
FtsI Speed (nm/s)
FtsI Histogram
Fast Pop
Slow Pop
